# Dual Oxygen-Partitioned Co-Culture Uncovers Microbe-Specific Epithelial Stress and Homeostatic Programs

**DOI:** 10.64898/2026.01.12.698921

**Authors:** Jae-Eun Lee, Taewon Kim, Seongwon Park, Ji-Young Lee, Kyoung Su Kim, Hong Koh, Keunwook Lee, Dong-Woo Lee, Nam Joo Kang

**Author notes:** Corresponding authors: Keunwook Lee, Department of Biomedical Science, Hallym University, Chuncheon 24252, Republic of Korea.; Dong-Woo Lee, Department of Biotechnology, Yonsei University, Seoul 03722, Republic of Korea.; Nam Joo Kang, Department of Biotechnology, The Catholic University of Korea, Bucheon 14662, Republic of Korea. These authors contributed equally to this work (Jae-Eun Lee, Taewon Kim, Seongwon Park).

## Abstract

Direct epithelial-microbial interactions occur across a steep aerobic-anaerobic interface in the intestine, yet mechanistic analysis has been limited by the difficulty of sustaining oxygen-dependent epithelial cells together with strictly anaerobic microbes *in vitro*. Here, we establish a dual oxygen-partitioned co-culture system that reproducibly maintains aerobic intestinal epithelial cells and obligate anaerobic gut bacteria, enabling controlled analysis of epithelial responses under physiologically relevant oxygen architecture. Using *Mediterraneibacter gnavus* and *Lacticaseibacillus casei* as contrasting microbial partners, we identify distinct epithelial programs. *M. gnavus* induces epithelial stress characterized by tight-junction disruption, suppression of mitochondrial respiration, and activation of chromatin-associated and innate immune regulatory pathways, whereas *L. casei* preserves barrier integrity and supports mitochondrial capacity with restrained immune modulation. Integrated transcriptomic and proteomic analyses reveal a shared epithelial energy-conserving response to microbial proximity, upon which microbe-specific stress or homeostatic programs are imposed. *In vivo* analysis using a dextran sodium sulfate-induced colitis model demonstrates that these epithelial programs become physiologically relevant under inflammatory conditions, with *L. casei* promoting epithelial repair and immune rebalancing. Together, these findings define oxygen-partitioned epithelial-microbial interactions as a determinant of microbe-specific epithelial states.

## Introduction

Inflammatory bowel disease (IBD), including ulcerative colitis (UC) and Crohn’s disease (CD), is a chronic inflammatory disorder of the gastrointestinal tract with a rapidly increasing global incidence (*1, 2*). A defining pathological feature of IBD is disruption of the intestinal epithelial barrier, which increases paracellular permeability and permits translocation of luminal antigens and microbial products (*3, 4*). Barrier failure initiates cytokine-driven inflammatory cascades, including tumor necrosis factor-α (TNF-α), interleukin-6 (IL-6), and IL-8, thereby amplifying mucosal immune dysregulation and disease progression (*5, 6*). These inflammatory responses are dynamically shaped by microbial composition, dietary cues, and epithelial metabolic state, underscoring the central role of epithelial-microbiome interactions in maintaining barrier integrity (*7, 8*). Beyond immune signaling, epithelial metabolic fitness, particularly mitochondrial function, has emerged as a critical determinant of barrier maintenance and repair. Mitochondrial dysfunction is increasingly observed in IBD epithelium (*9–12*) and is associated with impaired energy metabolism, defective barrier restoration, and failure to re-establish mucosal homeostasis (*13*).

Gut microbiota dysbiosis is a central feature of IBD and is characterized by loss of beneficial taxa, including *Lactobacillus*, *Bifidobacterium*, and other health-associated Bacillota (*14, 15*). Dysbiosis also perturbs microbial metabolite production, notably reducing short-chain fatty acids (SCFAs) that normally restrain TNF-α-driven inflammation and support epithelial energy metabolism (*16, 17*). Among disease-associated species, *Mediterraneibacter gnavus* is consistently enriched in UC and CD cohorts (*18, 19*). As a mucin-degrading anaerobe, *M. gnavus* erodes the mucus barrier and produce pro-inflammatory polysaccharides that activate dendritic cells and exacerbate epithelial stress (*20, 21*). By contrast, *Lacticaseibacillus casei* is a well-characterized probiotic shown to enhance barrier integrity, moderate cytokine signaling, and promote immune homeostasis in experimental colitis models (*22–24*). These opposing microbial phenotypes highlight the need for experimental systems that enable direct, controlled comparison of microbe-specific effects on epithelial physiology and metabolism.

Despite their close spatial proximity in the colon, epithelial-microbial interactions occur across a steep aerobic-anaerobic interface. Direct mechanistic analysis of these interactions has been limited by the difficulty of simultaneously maintaining oxygen-dependent epithelial cells and strictly anaerobic microbes *in vitro* (*25–27*). Existing platforms, including Transwell-based co-cultures, dynamic bioreactor systems such as the Simulator of the Human Intestinal Microbial Ecosystem (SHIME), and microfluidic devices such as gut-on-chip and the Human Microbial Cross-talk (HuMiX), have provided important insights into host-microbe interactions (*28–32*). However, these systems often require specialized instrumentation, lack precise control of vertical oxygen gradients, or do not support sustained epithelial–anaerobe contact. Organoid-based models more closely recapitulate epithelial architecture but remain poorly suited for long-term anaerobic co-culture (*33, 34*). Consequently, direct comparison of microbe-specific epithelial programs under defined oxygen-partitioned conditions has remained challenging.

To overcome these limitations, we optimized the human oxygen-bacteria anaerobic (HoxBan) co-culture system, originally developed for *Faecalibacterium prausnitzii* (*29*), to support robust and quantifiable co-culture of *M. gnavus* and *L. casei* with intestinal epithelial cells under a stable vertical oxygen gradient. Using this dual oxygen-partitioned system, we systematically investigated how contrasting gut microbes reshape epithelial barrier integrity, mitochondrial function, and immune-regulatory states through integrated transcriptomic and proteomic analyses, functional assays, and *in vivo* validation using a dextran sodium sulfate (DSS)-induced colitis model. Collectively, this work establishes an accessible and physiologically relevant platform for dissecting microbe-specific epithelial stress and homeostatic programs and provides mechanistic insight into how epithelial-microbial interactions become consequential under inflammatory conditions such as IBD.

## Methods

### Culture conditions

Human colorectal adenocarcinoma Caco-2 cells (ATCC HTB-37) were cultured in Eagle’s minimum essential medium (EMEM) supplemented with 20% fetal bovine serum (FBS) and 1% penicillin-streptomycin at 37°C. *Mediterraneibacter gnavus* ATCC 29149 and *Lacticaseibacillus casei* ATCC 393 were cultured in Tryptic soy broth (TSB), de Man, Rogosa, and Sharpe (MRS) medium under anaerobic conditions at 37°C. Gelrite-based matrices and mucin overlays were used to establish the oxic-anoxic co-culture interface. Antibodies for epithelial and tight junction marker analyses and reagents for mitochondrial respiration assays were obtained from commercial sources as specified in the **Supplementary Materials and Methods**.

### Dual oxygen (HoxBan) co-culture system

*M. gnavus* and *L. casei* were grown from single colonies in TSB or MRS medium, respectively, and subcultured into fresh medium at mid-exponential phase. Bacterial cultures harvested at exponential growth were used for co-culture experiments. Caco-2 cells were seeded at 3×10^4^ cells/cm^2^ on 0.1% gelatin-coated cover glasses and cultured for 48 h. Prior to co-culture, the medium was replaced with antibiotic-free EMEM supplemented with 10% FBS, and cells were equilibrated for 1 h.

All anaerobic components were prepared inside an anaerobic chamber. Briefly, 15 mL of TSB (for *M. gnavus*) or MRS (for *L. casei*) supplemented with Gelrite and 0.03% (w/v) MgCl_2_ was dispensed into 50-mL conical tubes. Gelrite concentrations were adjusted to 0.5% (w/v) in TSB and 0.7% (w/v) in MRS. The media were allowed to solidify to form the basal anaerobic layer. Exponentially growing bacteria were then mixed with semi-solid medium containing Gelrite [0.2% (w/v) in TSB for *M. gnavus* and 0.5% (w/v) in MRS for *L. casei*] at temperatures below 40°C and layered (10 mL per tube) onto the solidified base. A mucin overlay (0.1% mucin, 0.3% Gelrite, 0.03% MgCl_2_) was added and deoxygenated by injection of a mixed gas (10% H_2_, 10% CO_2_, and 80% N_2_) prior to solidification. The assembled anaerobic and mucin layers were transferred to an aerobic workbench, and cover glasses bearing confluent Caco-2 cells were placed cell-side down on the mucin surface. Each tube was filled with 10 mL of antibiotic-free EMEM containing 10% FBS, loosely capped, and incubated at 37°C in a humidified 5% CO_2_ atmosphere for 24 h to establish a stable dual-oxygen gradient supporting both epithelial cells and anaerobic bacteria.

### Measurement of oxygen profiles

Vertical oxygen concentration profiles within the co-culture system were measured using a Microx 4 oxygen sensor equipped with a PSt7 needle-type probe (PreSens, Germany). The probe was advanced in 1 mm increments from the apical epithelial medium through the mucin layer to the bacterial gel layer, and oxygen concentrations were recorded according to the manufacturer’s instructions.

### Cell viability assay

After 24 h of co-culture, Caco-2 cells were gently washed with warm phosphate-buffered saline (PBS) and detached using 0.25% trypsin-EDTA. Cell suspensions were collected in EMEM supplemented with 20% FBS and centrifuged at 3,000 rpm for 3 min. Pellets were resuspended in fresh medium and mixed 1:1 with 0.4% trypan blue. Viable (unstained) and non-viable (blue-stained) cells were quantified using a Countess II Automated Cell Counter (Thermo Fisher Scientific).

### Immunofluorescence microscopy

Immunofluorescence microscopy was used to assess epithelial identity, polarization, and tight junction integrity under conventional culture conditions and within the HoxBan system. Caco-2 cells cultured on cover glasses were fixed, permeabilized, and immunostained for epithelial markers, apical-basolateral polarity markers, or tight junction proteins using standard protocols. Confocal or fluorescence microscopy was performed to evaluate marker localization and epithelial architecture, and Z-stack imaging was used to assess epithelial polarization. Detailed antibody information, staining conditions, and imaging parameters are provided in the **Supplementary Materials and Methods**.

### Mitochondrial respiration analysis

Mitochondrial respiration in Caco-2 cells was assessed using the Seahorse XFp Cell Mito Stress Test. Cells recovered from the HoxBan system were replated in XFp miniplates and analyzed for oxygen consumption rate (OCR) following sequential inhibition of mitochondrial respiratory complexes. Basal respiration, ATP-linked respiration, and maximal respiratory capacity were calculated according to the manufacturer’s instructions. OCR values were normalized to viable cell numbers. Detailed assay conditions are described in the **Supplementary Materials and Methods**.

### Western blotting

Protein expression in Caco-2 cells and mouse colon tissues was analyzed by Western blotting. Total protein lysates were prepared following co-culture or tissue collection, separated by SDS-PAGE, and transferred to PVDF membranes. Membranes were probed with antibodies against epithelial junction proteins and detected using chemiluminescence. Detailed lysis conditions, antibody information, and detection methods are provided in the **Supplementary Materials and Methods**.

### Transcriptome analysis

Total RNA was extracted from Caco-2 cells following co-culture with or without gut bacteria in the HoxBan system and subjected to RNA sequencing. Differential gene expression analysis was performed to identify epithelial transcriptional responses to microbial co-culture. Principal component analysis was used to assess global transcriptional variation between conditions. Functional enrichment analyses were conducted to characterize stress-, immune-, and metabolism-related pathways, including targeted evaluation of mitochondrial-associated gene expression. Detailed sequencing procedures, data processing, and statistical analyses are described in **the Supplementary Materials and Methods**.

### Proteomic analysis

Proteomic profiling of Caco-2 cells, co-cultured bacteria, and mouse colon tissues was performed using liquid chromatography-tandem mass spectrometry (LC-MS/MS). Proteins were extracted, enzymatically digested, and analyzed by high-resolution Orbitrap mass spectrometery. Label-free quantification was used to identify differentially expressed proteins in response to microbial co-culture or in vivo bacterial transfer. Functional enrichment analyses were conducted to characterize metabolic, immune, and stress-related pathways. Detailed sample preparation, mass spectrometry settings, database searches, and statistical criteria are described in the **Supplementary Materials and Methods**.

### Cytokine array assay

Conditioned media from the apical compartment of the HoxBan co-culture systems were applied to human peripheral blood mononuclear cells (hPBMCs), and cytokine secretion profiles were analyzed using a membrane-based human cytokine array. Signal intensities were quantified and normalized to internal controls to compare immune activation patterns induced by different microbial conditions.

Detailed experimental procedures and data normalization methods are described in the **Supplementary Materials and Methods**.

### DSS-induced colitis mouse model

A dextran sodium sulfate (DSS)-induced colitis model was used to evaluate *in vivo* relevance of HoxBan-derived host-microbe interactions. Antibiotic-pretreated C57BL/6J mice were orally administered *M. gnavus* or *L. casei* prior to DSS exposure (**Fig. 5A**). Disease activity, epithelial integrity, and inflammatory responses were assessed using established clinical and molecular readouts. Full animal handling, treatment regimens, and scoring criteria are provided in the **Supplementary Materials and Methods**.

### Histopathology, immunofluorescence, and immune profiling

Distal colon tissues were subjected to histopathological evaluation, immunofluorescence staining, and immune profiling to assess epithelial integrity, inflammatory status, and immune cell composition following bacterial transfer in DSS-induced colitis. Goblet cell preservation, epithelial proliferation, mitochondrial content, cytokine production, and immune cell populations in the colonic lamina propria were analyzed using established staining, flow cytometry, and multiplex cytokine assays. Gut microbiota composition was assessed by 16S rRNA gene amplicon sequencing. Detailed experimental procedures and analytical pipelines are provided in the **Supplementary Materials and Methods**.

### Statistical analysis

Data are presented as mean ± standard deviation (SD). For comparisons between two groups, two-tailed Student’s *t*-tests were used. For *in vivo* experiments, statistical significance was assessed relative to healthy or DSS control groups as indicated in the figure legends. Differences were considered statistically significant at *p* < 0.05. The number of biological replicates (n) and exact statistical comparisons are specified in the figure legends.

## Results

### Development of an oxic-anoxic co-culture system defining a biologically permissive window for epithelial-microbial interactions

Although the original HoxBan system enabled aerobic-anaerobic epithelial-microbial co-culture, it did not define the biologically permissive conditions required for mechanistic interpretation, owing to limited visualization of bacterial growth and insufficient validation of oxygen distribution across culture layers (*29*). Without such definition, direct microbial effects are difficult to distinguish from nonspecific epithelial stress. To address this limitation, we introduced three key modifications: (1) replacement of agar with a transparent Gelrite matrix to enable direct visualization of bacterial expansion; (2) incorporation of a mucus-like gel layer to approximate the intestinal barrier; and (3) quantitative oxygen profiling to validate establishment of a physiologically relevant vertical oxygen gradient. The optimized system comprises a semi-solid bacterial matrix, a mucin-coated interface, an inverted epithelial monolayer, and an aerobic liquid compartment, collectively recreating the oxic-anoxic architecture of the intestinal lumen (**Fig. 1A–B**).

**Figure 1.**
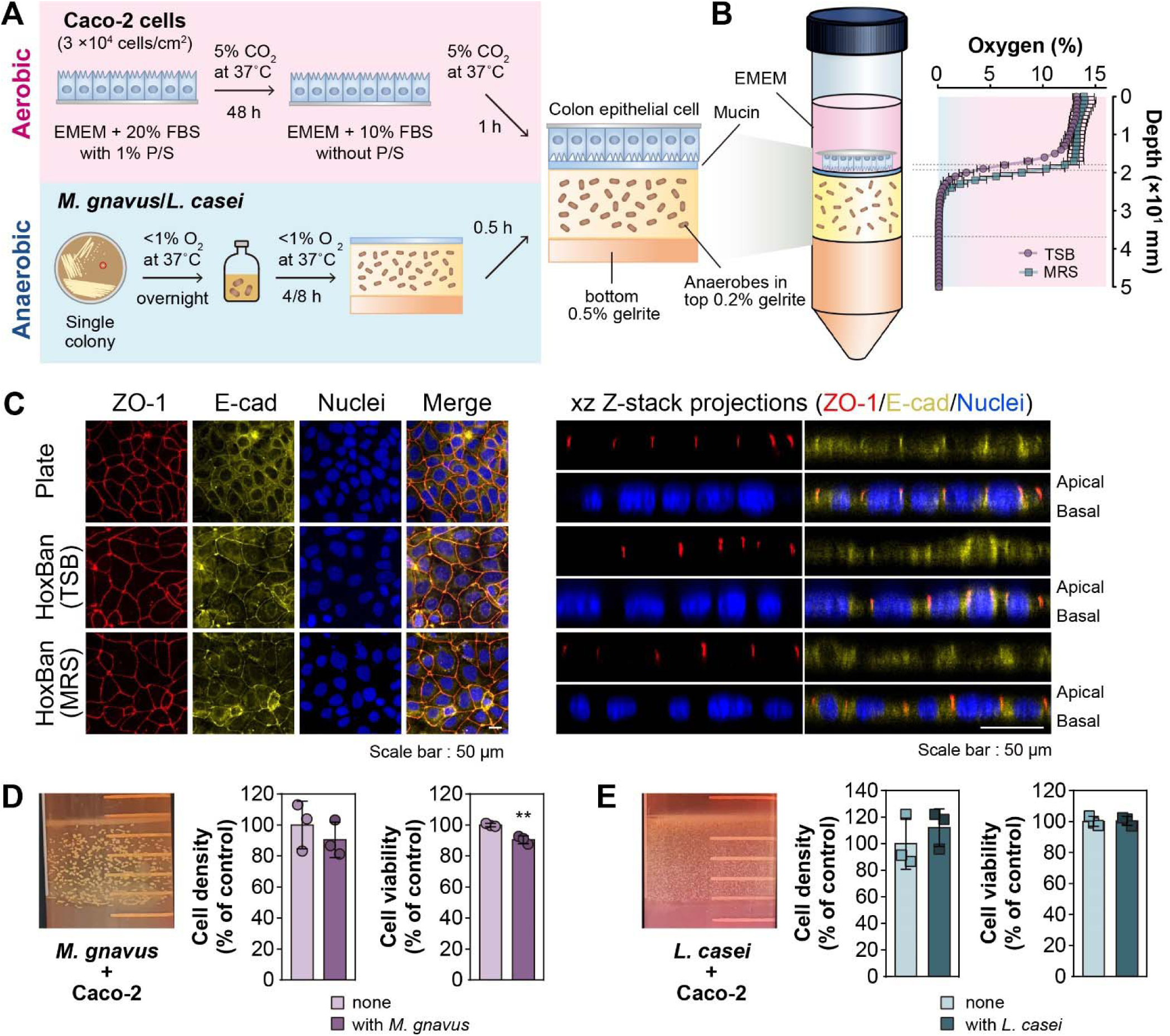
Establishment of an oxic-anoxic co-culture system supporting epithelial and anaerobic microbial growth. **(A)** Schematic overview of preparation of aerobic epithelial cells and anaerobic gut bacteria prior to assembly of the HoxBan co-culture system. **(B)** Schematic representation of the assembled HoxBan system and the vertical oxygen gradient across culture layers. Oxygen concentrations were measured from the apical epithelial medium to the basal bacterial matrix using a needle-type oxygen probe. Dashed lines indicate layer interfaces. **(C)** Confocal z-stack imaging of Caco-2 cells cultured on conventional plates or in the HoxBan system (TSB or MRS) in the absence of bacteria. ZO-1 (red), E-cadherin (yellow), and nuclei (DAPI, blue) are shown. Representative xy and xz projections demonstrate preserved epithelial organization under oxic-anoxic conditions. **(D)** Representative image of *M. gnavus* growth in the HoxBan system after 24 h of co-culture (initial inoculum: 5×10² cells/tube), with corresponding Caco-2 cells density and viability assessed by Trypan Blue exclusion. **(E)** Representative image of *L. casei* growth after 24 h co-culture (initial inoculum: 1×10□ cells/tube), with corresponding epithelial cell density and viability. Quantitative data are presented as the mean ± SD (n=3). Statistical significance was determined by two-tailed Student’s *t*-test compared with the corresponding bacteria-free condition (** *p* < 0.01). Scale bars, 50 µm.

*M. gnavus* and *L. casei* were activated under strict anaerobic conditions and introduced into semi-solid media during exponential growth (**Fig. 1A**). Caco-2 cells cultured on gelatin-coated coverslips were grown to approximately 80-90% confluence and inverted onto the mucin-coated microbial layer. Co-cultures were maintained for 24 h under defined oxygen-gradient conditions. Direct oxygen profiling confirmed a stable vertical gradient, with ∼15% O_2_ in the upper liquid phase, 10–12% at the epithelial-mucin interface, and <1% within the bacterial matrix (**Fig. 1B**). Although these absolute oxygen levels exceed those of the distal colon, preservation of a directional gradient across the epithelial-luminal axis provides the dominant physiological cue governing epithelial-microbial interactions.

We next assessed whether epithelial baseline properties were preserved within this permissive window. Confocal differential interference contrast (DIC) imaging revealed intact cobblestone morphology comparable to conventional plate cultures (**Fig. S1A–B**). Core epithelial markers, including the tight junction protein ZO-1 (*35*), adherens junction protein E-cadherin (*36*), and the cytoskeletal marker pan-cytokeratin (*37*), were maintained. Mitochondrial respiration, assessed by oxygen consumption rate (OCR), showed no significant differences between HoxBan and standard culture conditions regardless of bacterial medium composition, indicating preserved epithelial metabolic competence (**Fig. S1C–E**). Confocal z-stack imaging further demonstrated preserved epithelial organization, with ZO-1 localized to intracellular junctions and E-cadherin confined to lateral membranes (**Fig. 1C**), supporting maintenance of polarized epithelial architecture.

To define microbial conditions compatible with epithelial viability, we next optimized bacterial inoculum density. *M. gnavus* induced inoculum-dependent epithelial stress, with marked loss of viability at ≥10^3^ cells/tube, whereas an inoculum of 5 × 10^2^ cells/tube preserved ∼87% epithelial viability and was selected for subsequent experiments (**Figs. 1D, S2A, and S2B**). In contrast, *L. casei* maintained epithelial viability across all tested inocula, and an inoculum of 10^5^ cells/tube was selected as the working condition (**Figs. 1E, S2C, and S2D**).

Together, these results establish an oxic-anoxic co-culture system that defines a biologically permissive window in which epithelial integrity, metabolic function, and oxygen architecture are preserved while enabling controlled interaction with anaerobic gut microbes. This defined operating window provides a critical foundation for attributing downstream epithelial phenotypes to microbe-specific activities rather than nonspecific cytotoxicity.

### Co-culture with *M. gnavus* and *L. casei* exerts opposing effects on tight junction integrity and mitochondrial function in Caco-2 cells

Using the optimized HoxBan platform, we examined how direct co-culture with anaerobic gut microbes alters epithelial functions central to intestinal homeostasis, focusing on tight junction (TJ) integrity and mitochondrial respiration. Tight junctions are essential for maintaining epithelial barrier function, and their disruption facilitates luminal antigen translocation and inflammatory activation (*38, 39*). In Caco-2 cells co-cultured with *M. gnavus*, immunofluorescence microscopy revealed markedly reduced junction-associated localization of TJ proteins, including ZO-1 and occludin, accompanied by a significant decrease in total protein abundance as determined by Western blotting (**Fig. 2A–B**). In contrast, co-culture with *L. casei* preserved and enhanced TJ integrity, as evidenced by stronger ZO-1 and occludin signals together with increased protein abundance detected by Western blotting (**Fig. 2C–D**). These findings demonstrate pronounced, species-dependent effects on epithelial barrier integrity.

**Figure 2.**
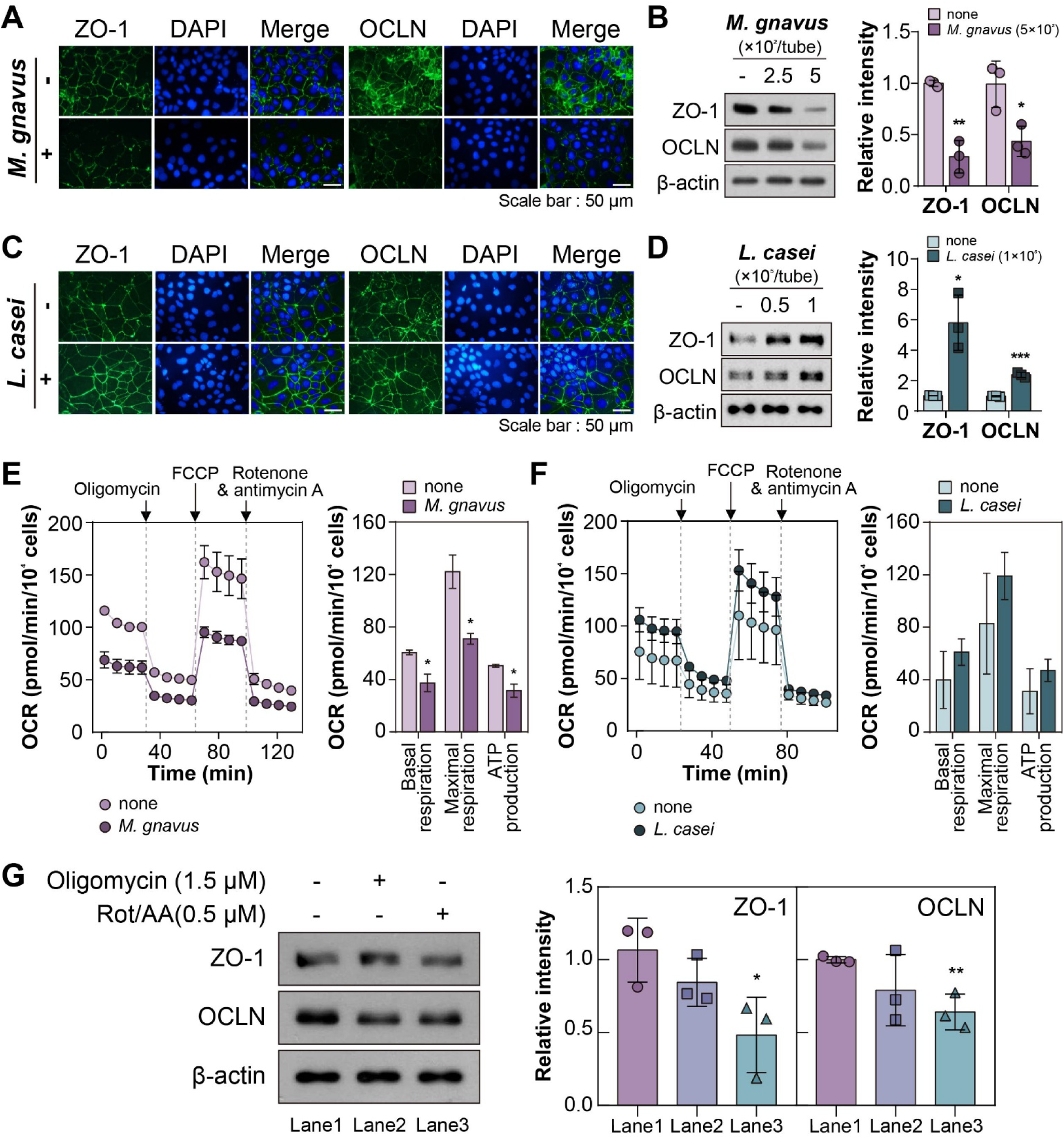
Opposing effects of gut bacteria on epithelial tight junction integrity and mitochondrial respiration. **(A)** Immunofluorescence (IF) staining of ZO-1 and occludin (OCLN) in Caco-2 cells after 24 h co-culture with *M. gnavus*. Nuclei were counterstained with DAPI. **(B)** Western blot analysis and quantification of ZO-1 and OCLN following *M. gnavus* co-culture, normalized to β-actin and expressed relative to bacteria-free controls. **(C)** IF staining of ZO-1 and OCLN after co-culture with *L. casei*. **(D)** Western blot analysis and quantification of ZO-1 and OCLN following *L. casei* co-culture. **(E)** Representative oxygen-consumption rate (OCR) traces of Caco-2 cells after *M. gnavus* co-culture measured using a Seahorse XFp analyzer. **(F)** OCR traces and derived mitochondrial respiration parameters following *L. casei* co-culture. **(G)** Effects of mitochondrial respiratory inhibition on TJ protein expression. Caco-2 cells were treated with oligomycin or rotenone/antimycin A under standard cuture conditions, followed by Western blot analysis. Data represent three independent experiments and are shown as mean ± SD (n = 3). \**p* < 0.05; \*\**p* < 0.01; \*\*\**p* < 0.001.

We next assessed whether bacterial co-culture modulates epithelial bioenergetics. OCR analysis showed that *M. gnavus* significantly suppressed mitochondrial respiration, reducing basal respiration, maximal respiratory capacity, and ATP-linked oxygen consumption by 38.3%, 42.0%, and 37.9%, respectively (**Fig. 2E**). In contrast, *L. casei* maintained mitochondrial respiratory activity at levels comparable to control cultures, with a modest upward trend across OCR parameters (**Fig. 2F**). To determine whether impaired mitochondrial respiration directly contributes to TJ stability, mitochondrial function was independently inhibited under standard culture conditions. Inhibition of ATP synthase with oligomycin alone did not substantially alter TJ protein expression, whereas blockade of electron transport using rotenone/antimycin A resulted in a clear reduction of both ZO-1 and occludin levels (**Fig. 2G**). These results indicate that disruption of mitochondrial electron transport is sufficient to compromise epithelial barrier integrity.

Together, these data demonstrate that *M. gnavus* weakens epithelial barrier structure in parallel with suppression of mitochondrial respiration, whereas *L. casei* preserves TJ integrity while maintaining epithelial bioenergetic function. These opposing epithelial responses highlight the utility of the oxic-anoxic co-culture system for resolving species-dependent host–microbe interactions under physiologically relevant conditions.

### Gut bacteria remodel epithelial transcriptomes through distinct stress- and immune-regulatory programs

To define epithelial responses beyond alterations in TJ integrity and mitochondrial function, we profiled transcriptomes of Caco-2 cells co-cultured with *M. gnavus* or *L. casei* using the optimized HoxBan system. RNA-seq libraries exhibited high sequencing quality, with >98% of reads mapping to the human genome (**Table S1**). Principal component analysis (PCA) revealed clear separation of both co-culture conditions from bacteria-free controls, indicating broad transcriptional remodeling induced by microbial exposure (**Fig. 3A**). Differential expression analysis identified 443 upregulated and 616 downregulated transcripts in response to *M. gnavus*, whereas *L. casei* induced a more moderate response with 201 upregulated and 479 downregulated transcripts (**Fig. 3B; Dataset 1**). When restricted to protein-coding genes, *M. gnavus* elicited substantially larger transcriptional changes (201 upregulated and 479 downregulated genes) than *L. casei* (69 upregulated and 253 downregulated genes), consistent with the stronger functional phenotypes observed at the epithelial level.

**Figure 3.**
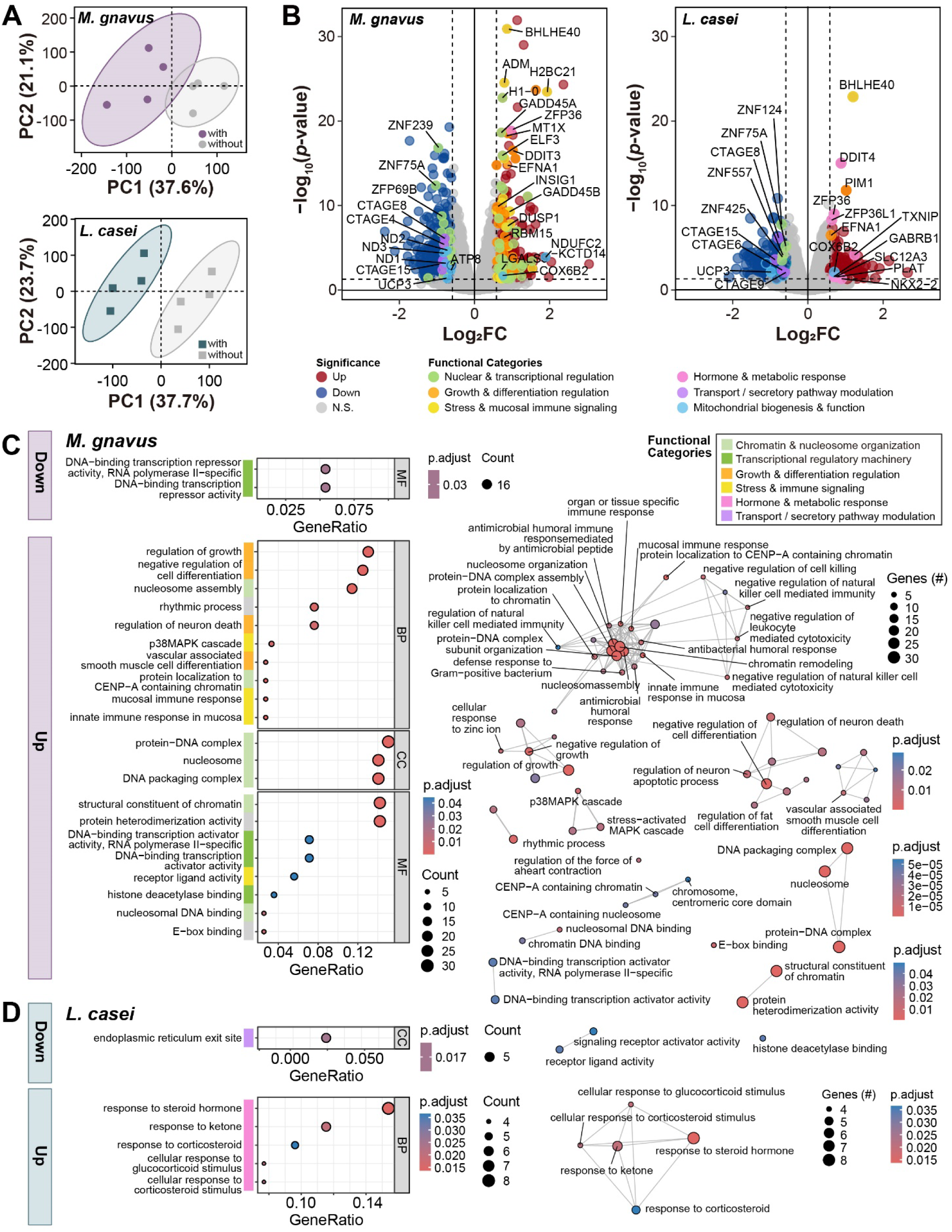
Transcriptomic divergence of epithelial states induced by anaerobic gut microbes. (**A**) Principal component analysis (PCA) of RNA-seq profiles from Caco-2 cells after 24 h co-culture with *M. gnavus* (top) or *L. casei* (bottom). Shaded ellipses indicate 95% confidence intervals. (**B**) Volcano plots showing differentially expressed genes (DEGs) following co-culture with *M. gnavus* (left) or *L. casei* (right). Upregulated and downregulated genes are shown in red and blue, respectively. Representative DEGs are annotated by functional categories. (**C**−**D**) Gene ontology (GO) enrichment analysis of DEGs induced by microbial co-culture (*M. gnavus,* **C**; *L. casei,* **D**). Dot plots and enrichment maps summarize significantly enriched Biological Process (BP); Cellular Component (CC), and Molecular Function (MF) terms for downregulated or upregulated genes. Dot plots display the top 10 enriched terms, and enrichment maps summarize all significant terms (adjusted *p* < 0.05). GO terms are grouped into functional categories as indicated. Full enrichment results are provided in **Dataset 2**.

Gene ontology (GO) analysis revealed pronounced reorganization of epithelial transcriptional regulation in *M. gnavus* co-cultures (**Fig. 3B–C; Dataset 2**). Downregulated genes were significantly enriched for DNA-binding transcription repressor activity (GO:0001217, FDR = 0.0296), driven by coordinated repression of multiple zinc-finger (ZNF) family members, including ZNF75A, ZNF79, ZFP69B, ZNF180, and ZNF239 (**Fig. 3C; Dataset 2**). In contrast, upregulated genes were enriched for transcription activator activity (GO:0001216; FDR = 0.0409; GO:0001228; FDR = 0.0409), indicating a global shift in transcriptional control.

The dominant upregulated signature in *M. gnavus*-exposed cells comprised chromatin- and nucleosome-associated programs, including structural constituent of chromatin (GO:0030527, FDR = 1.0 × 10 ^31^), nucleosome (GO:0000786, FDR = 1.5 × 10 ^26^), DNA packaging complex (GO:0044815, FDR = 3.2 × 10 ^23^), and nucleosome assembly (GO:0006334, FDR = 2.9 × 10 ^17^) (**Fig. 3C; Dataset 2**). Mucosa-associated innate immune categories, including innate immune response in mucosa (GO:0002227, FDR = 3.5 × 10 ^4^) and mucosal immune response (GO:0002385, FDR = 1.9 × 10 ^3^), were also enriched, but were primarily supported by histone-associated genes (e.g., H2BC8, H2BC7, H2BC4, H2BC21, and H2BC11) rather than classical immune effector genes. Consistent with stress-responsive signaling, *M. gnavus* exposure additionally enriched regulation of the p38 MAPK cascade (GO:0038066, FDR = 4.9 × 10 ^3^).

Given the pronounced mitochondrial phenotype observed functionally, mitochondrial-associated genes were examined using targeted annotation. This analysis revealed coordinated downregulation of multiple electron transport chain (ETC) components, including Complex I subunits (ND1, ND2, ND3) and ATP synthase subunit ATP8, with limited compensatory induction of other mitochondrial transcripts (**Fig. 3B; Dataset 1)**. Although these changes did not form an enrichment-defined mitochondrial GO category, their coordinated nature is consistent with the observed suppression of oxygen consumption (**Fig. 2E**). Integrated gene-level and gene set-level analyses thus define a *M. gnavus*-specific transcriptional signature characterized by induction of stress- and immune-regulatory programs alongside repression of mitochondrial ETC, a pattern not observed in *L. casei* co-cultures (**Fig. 3B–D**).

Comparison with immune reference gene sets from MSigDB (C7) revealed significant overlap between *M. gnavus*–regulated genes and transcriptional signatures derived from LPS- or TLR-stimulated dendritic cells and macrophages, as well as activated CD4^+^ T cells (**Fig. S3A–B; Dataset 2**). These overlaps were driven primarily by stress-responsive regulators, including GADD45A, DUSP1 and PER1, indicating engagement of immune-regulatory stress programs rather than lineage-specific immune differentiation.

In contrast, *L. casei* induced a comparatively restrained transcriptional response. The only significantly enriched downregulated category was endoplasmic reticulum exit site (GO:0070971, FDR = 0.0168) (**Fig. 3D; Dataset 2**). Genes within this category showed concordant downregulation under both co-culture conditions, suggesting a shared epithelial response to microbial proximity rather than a species-specific effect (**Fig. 3B)**. Upregulated genes in *L. casei* co-cultures were primarily enriched for hormone- and metabolism-related pathways (GO:0048545, GO:0071385, GO:1901654; FDR < 0.02), with minimal immune-associated enrichment (**Figs. 3D and S3C; Dataset 2**).

Collectively, these transcriptomic analyses demonstrate that *M. gnavus* drives a stress-adaptive, chromatin-remodeling epithelial program coupled to partial repression of mitochondrial electron transport, whereas *L. casei* elicits a more limited epithelial transcriptional response biased toward metabolic and hormonal regulation. These findings highlight how distinct gut microbes induce fundamentally different epithelial gene-expression programs and highlight the utility of the HoxBan system for resolving microbe-dependent epithelial reprogramming under physiologically relevant oxic-anoxic conditions.

### Proteomic profiling reveals energy-conserving and microbe-specific adaptive programs in co-cultured epithelial cells

To directly assess functional adaptations during epithelial−microbial co-culture, we independently profiled epithelial and bacterial proteomes recovered from the HoxBan system. Host-microbe separation was achieved by sequential trypsinization and alcalase digestion, enabling compartment-resolved LC-MS/MS analysis (**Figs. S4A–B and S5A–B**). Label-free quantitative proteomics identified 3,041 human proteins, ∼85% of which were consistently detected across all conditions (**Fig. S4C**). Despite this shared proteomic core, unsupervised clustering revealed clear segregation of epithelial proteomes according to bacterial exposure **(Fig. S4D**), indicating robust microbe-dependent remodeling. In Caco-2 cells, co-culture with *M. gnavus* resulted in 65 significantly upregulated and 172 significantly downregulated proteins, whereas *L. casei* induced 85 upregulated and 185 downregulated proteins **(Fig. 4A; Dataset 3**). Across both microbial conditions, epithelial cells exhibited a conserved adaptative response characterized by suppression of energy-intensive biosynthetic processes, including protein synthesis and cell-cycle-associated functions **(Figs. 4B–C; Dataset 4).** This shared response is consistent with a generalized energy-conserving state during bacterial exposure.

**Figure 4.**
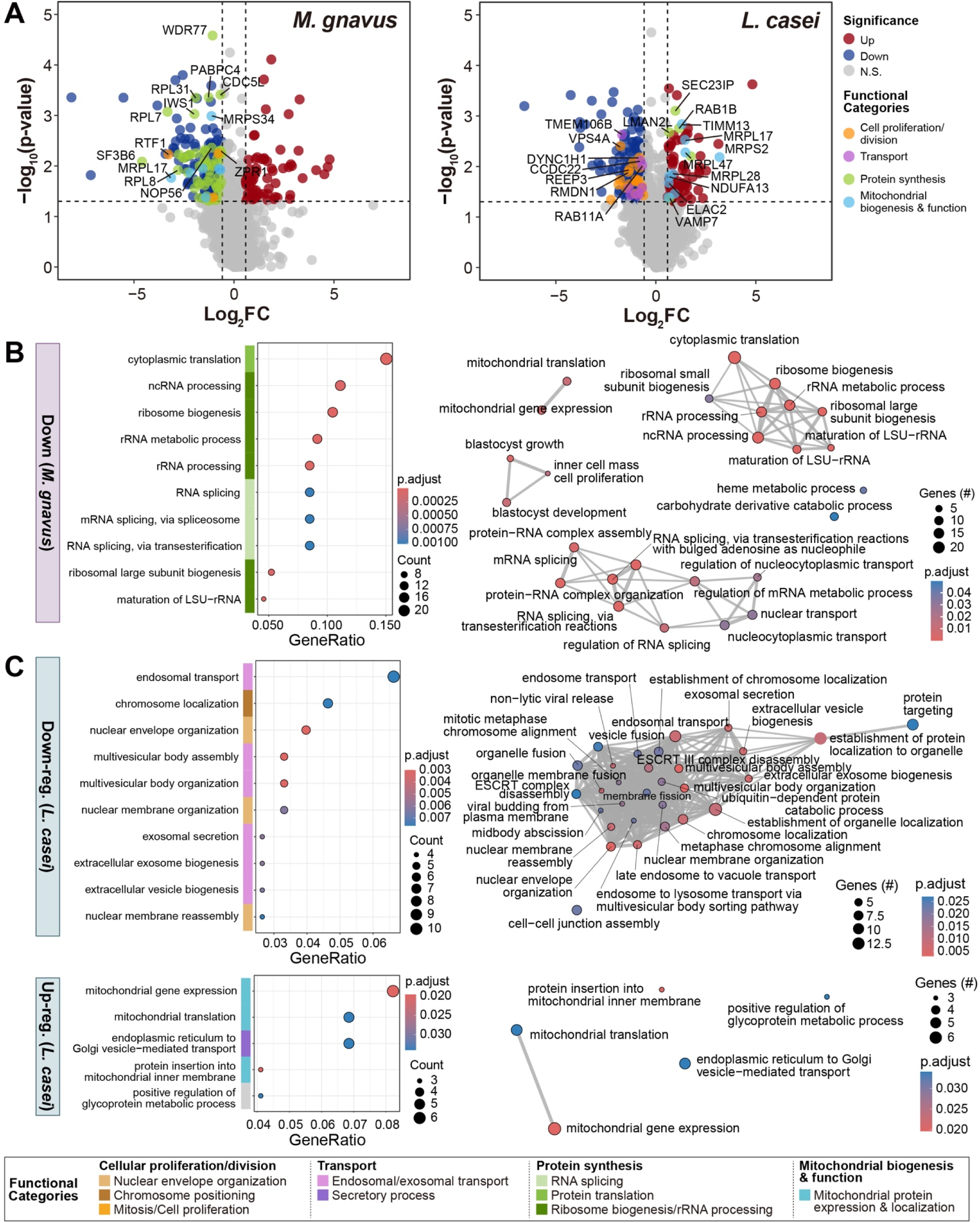
Proteomic remodeling of epithelial cells during microbial co-culture. (**A**) Volcano plots of differentially expressed proteins in Caco-2 cells after co-culture with *M. gnavus* (left) and *L. casei* (right). Proteins involved in cell proliferation/division, transport, protein synthesis, and mitochondrial biogenesis & function are highlighted. (**B**−**C**) GO enrichment analysis of differentially expressed proteins following co-culture with *M. gnavus* (**B**) or *L. casei* (**C**). Dot plots show the top enriched terms, and enrichment maps summarize all significant categories (adjusted *p* < 0.05). Full enrichment results are provided in **Dataset 4**.

Notably, the regulatory routes underlying this conserved phenotype differed markedly between the two microbes. In *M. gnavus* co-cultures, downregulated proteins were strongly enriched for translational and RNA-processing pathways, including ribosome biogenesis (GO:0042254, FDR = 7.4 × 10^-6^), RNA processing (GO:0006364, FDR = 1.6 × 10^-5^; GO:0034470, FDR = 3.8 × 10^-5^), and translation-related processes (e.g., GO:0002181, FDR =4.4 × 10^-19^) (**Fig. 4A–B**). In contrast, repression of cell-division-associated processes was more prominent in *L. casei* co-cultures, including nuclear envelope organization (GO:0006998, FDR = 3.0 × 10^-3^), chromosome localization (GO:0050000, FDR = 7.6 × 10^-3^), and endosomal transport (GO:0016197, FDR = 7.6 × 10^-3^) (**Fig. 4C**). These data indicate that epithelial energy conservation is achieved primarily through translational suppression in response to *M. gnavus*, but through mitotic restraint in response to *L. casei*.

Species-specific mitochondrial adaptations were also evident at the proteomic level. Co-culture with *L. casei* selectively upregulated proteins involved in mitochondrial translation and import, including mitochondrial ribosomal proteins (MRPS35, MRPL28, MRPL17, MRPL47, and MRPS2) and TIM complex components (TIMM10B and TIMM13) (GO:0140053, FDR = 2.0 × 10^-2^; GO:0032543, FDR = 3.4 × 10^-2^; GO:0045039, FDR = 2.0 × 10^-2^) (**Fig. 4A** and **C**). Although most individual OXPHOS subunits were not strongly induced (e.g., NDUFA13, log_2_FC = 0.68, *p* = 0.016), the coordinated enhancement of mitochondrial translational and import capacity is consistent with the preserved or modestly enhanced OCR observed functionally (**Figs. 2F and 4A**). In contrast, *M. gnavus* co-culture led to coordinated downregulation of mitochondrial protein synthesis, in parallel with its broader suppression of cytosolic translation (**Fig. 4B**).

Immune-associated proteomic signatures further distinguished the two microbial conditions. Proteins downregulated in *M. gnavus* co-cultures significantly overlapped with infection-responsive immune signatures derived from bacterial, parasitic, and viral challenge models (**Fig. S6A; Dataset 4**), consistent with a stress-associated epithelial state. In contrast, *L. casei* co-cultures were enriched for cytokine-modulated innate and adaptive immune-related signatures, alongside attenuation of infection-associated protein patterns (**Fig. S6B; Dataset 4**), indicating a more regulated immune-modulatory epithelial response.

We next examined bacterial proteomes to identify microbial adaptations accompanying host contact. While the two species expressed largely overlapping protein sets, *M. gnavus* exhibited a stronger host-induced proteomic shift, with 180 proteins significantly upregulated and 161 downregulated in the presence of epithelial cells **(Fig. S5A; Dataset 3)**. Differentially expressed bacterial proteins were enriched for pathways related to cell-envelope remodeling, transcriptional regulation, and nutrient utilization **(Fig. S5C–D**). Notably, the two species exhibited opposing stress-response patterns: *M. gnavus* upregulated multiple DNA repair and stress-response proteins, whereas *L. casei* showed a general reduction in these categories. In addition, *M. gnavus* increased expression of mucin-degrading enzymes and pathways involved in sialic acid and fucose utilization, consistent with preferential consumption of mucin-derived substrates upon host association (*17*).

To determine whether bacteria-induced epithelial states translated into functional immune cues, conditioned media from co-cultures were applied to human PBMCs and analyzed using cytokine arrays (**Fig. S7A–B**). After normalization to controls and LPS responses (control = 0, LPS = 100), *M. gnavus* co-culture media induced strong pro-inflammatory cytokines, including IL-6 and IL-8, reaching levels comparable to LPS stimulation (**Fig. S7C–E; Dataset 5**). In contrast, *L. casei* media elicited only a modest increase in IL-8 and reduced IL-6, while preferentially elevating cytokines such as IL-3, IL-5, and IL-13, which are associated with immune modulation and epithelial remodeling rather than acute pro-inflammation responses.

Overall, integrated proteomic and cytokine analyses indicate that epithelial cells adopt a shared energy-conserving state during bacterial exposure, but engage distinct adaptive programs depending on microbial identity. *M. gnavus* drives a stress-responsive epithelial state characterized by translational and mitochondrial restraint and a pro-inflammatory immune tone, whereas *L. casei* promotes a restrained, immunomodulatory profile accompanied by enhanced mitochondrial capacity. These findings further underscore the utility of the HoxBan system for resolving microbe-dependent epithelial adaptations under physiologically relevant oxic–anoxic conditions.

### *In vivo* validation of HoxBan-derived host–microbe interactions in DSS-induced colitis

The optimized HoxBan system revealed that *M. gnavus* and *L. casei* elicit contrasting epithelial responses *in vitro*. *M. gnavus* induced junctional disruption, mitochondrial dysfunction, and stress-associated transcriptional programs, whereas *L. casei* enhanced barrier integrity, supported mitochondrial capacity, and exerted restrained immune modulation. To determine whether these epithelial phenotypes translate *in vivo*, we evaluated the effects of each bacterium in a dextran sulfate sodium (DSS)-induced colitis model. Mice were pretreated with amoxicillin to facilitate microbial engraftment (*40*), followed by oral administration of *M. gnavus* or *L. casei* prior to and during DSS exposure (**Fig. 5A**). DSS treatment induced characteristic colitis manifestations, including progressive body weight loss and elevated disease activity scores (**Fig. 5B–C**). Administration of *L. casei*, but not *M. gnavus*, significantly attenuated weight loss and reduced disease severity. Consistent with these clinical improvements, *L. casei* treatment decreased fecal inflammatory biomarkers, including lipocalin-2 and calprotectin (**Fig. S8A**), and partially restored colon length (**Fig. 5D**). Histopathological analysis further demonstrated that *L. casei* protected against DSS-induced epithelial damage, inflammatory cell infiltration, and goblet cell depletion, all hallmark features of ulcerative colitis (**Figs. 5E and S8B**). In contrast, *M. gnavus* administration did not significantly ameliorate histopathological injury.

**Figure 5.**
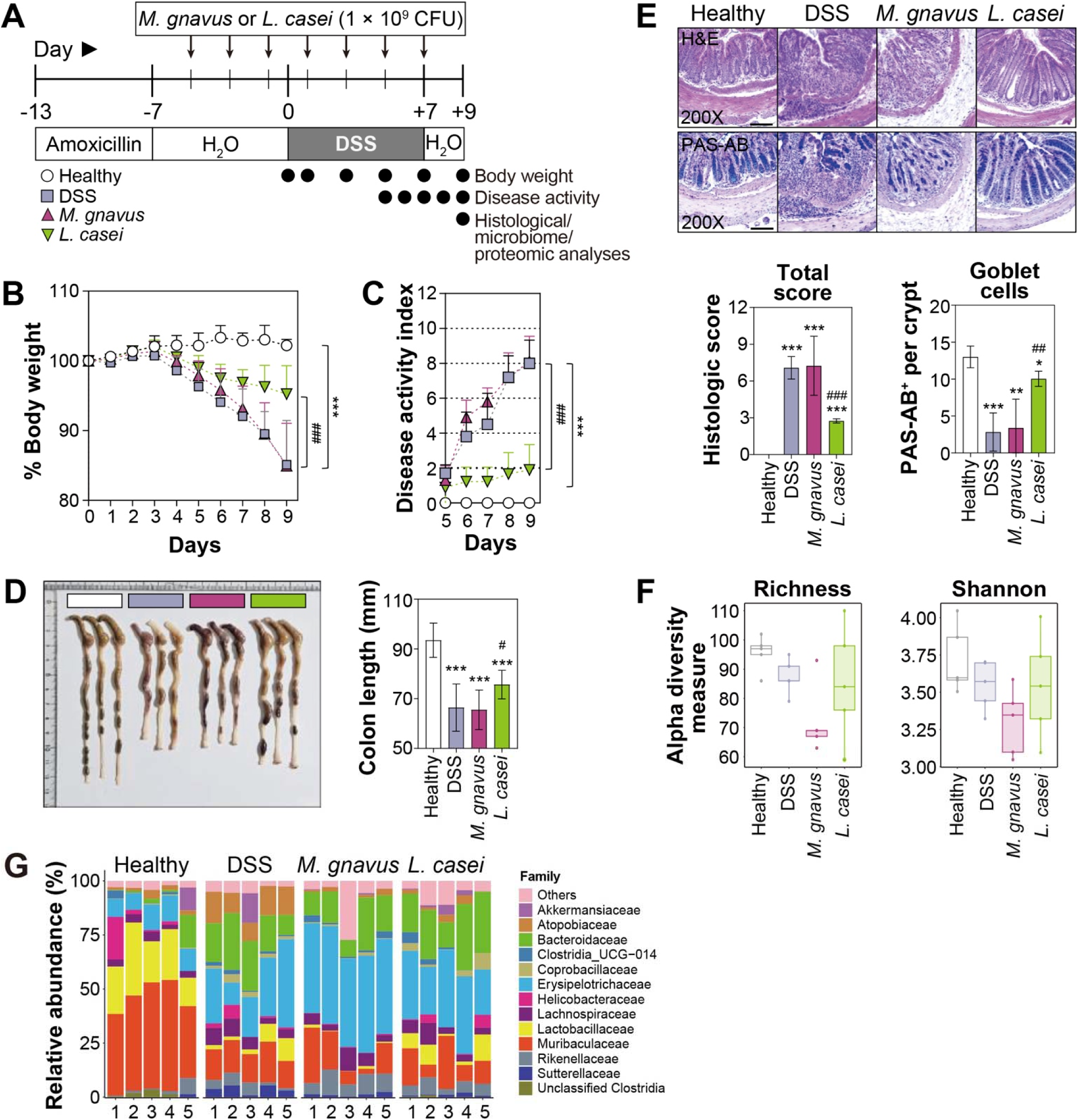
*In vivo* validation of host-microbe interactions in DSS-induced colitis. (**A**) Experimental timeline of DSS-induced colitis with microbial administration. After amoxicillin pretreatment, mice were orally administered *M. gnavus*, *L. casei*, or PBS (DSS control), followed by 2.5% DSS in drinking water to induce colitis (n = 10 per group). (**B** and **C**) Body weight change and disease activity index following DSS treatment. Statistical significance was determined relative to the healthy and DSS-induced groups (\*\*\**p* < 0.001 vs. healthy control; ^###^*p* < 0.001 vs. DSS control). (**D**) Colon length measured at day 9 after DSS treatment. (**E**) Representative histological images of distal colon stained with H&E and PAS–Alcian blue (200×: scale bar = 100 µm). Histological scores and goblet cell count per crypt were quantified and presented as bar graphs (n = 4). (**F**) Alpha-diversity indices (Richness and Shannon index) of fecal microbiota in the healthy, DSS-induced, *M. gnavus*–treated, and *L. casei*–treated groups (n = 5). (**G**) Family-level relative abundance of gut microbiota, with “Others” representing taxa below 1% relative abundance.

Because microbial supplementation can influence both host inflammation and the resident microbial community, we next assessed changes in gut microbiota composition. Although neither *M. gnavus* nor *L. casei* showed clear evidence of stable colonization, both altered microbial community structure under colitic conditions. Alpha diversity was highest in healthy mice, intermediate in *L. casei*-treated animals, and lowest in the *M. gnavus* group (**Fig. 5F**). NMDS ordination and compositional profiling revealed distinct microbiota restructuring in both treatment groups relative to DSS controls (**Fig. S8C–D**). At the family level, healthy mice were enriched in *Muribaculaceae* and *Lactobacillaceae* and showed reduced *Bacteriodaceae* abundance (**Fig. 5G**). LEfSe analysis further identified enrichment of *Ileibacterium*, *Rikenellaceae*_RC9_gut_group, and *Romboutsia* in the *M. gnavus* group, whereas *Turicibacter* was preferentially enriched following *L. casei* administration (**Fig. S8E**). Consistent with these compositional shifts, proteomic profiling of colon tissues revealed species-specific remodeling, with *L. casei*-treated mice showing the largest deviation from DSS-only proteomic profiles (**Fig. S8F-G**).

We next examined epithelial integrity and immune activation within the colon. DSS treatment markedly reduced expression of the TJ proteins occludin and claudin-1, whereas *L. casei* administration restored their expression toward healthy levels (**Fig. 6A**). In parallel, *L. casei* increased the abundance of Ki-67^+^ proliferating enterocytes and elevated epithelial mitochondrial mass (**Fig. 6B**), consistent with the mitochondrial-supportive and epithelial-repair phenotypes observed *in vitro*. In contrast, *M. gnavu*s administration did not significantly alter epithelial junctional markers or mitochondrial parameters.

**Figure 6.**
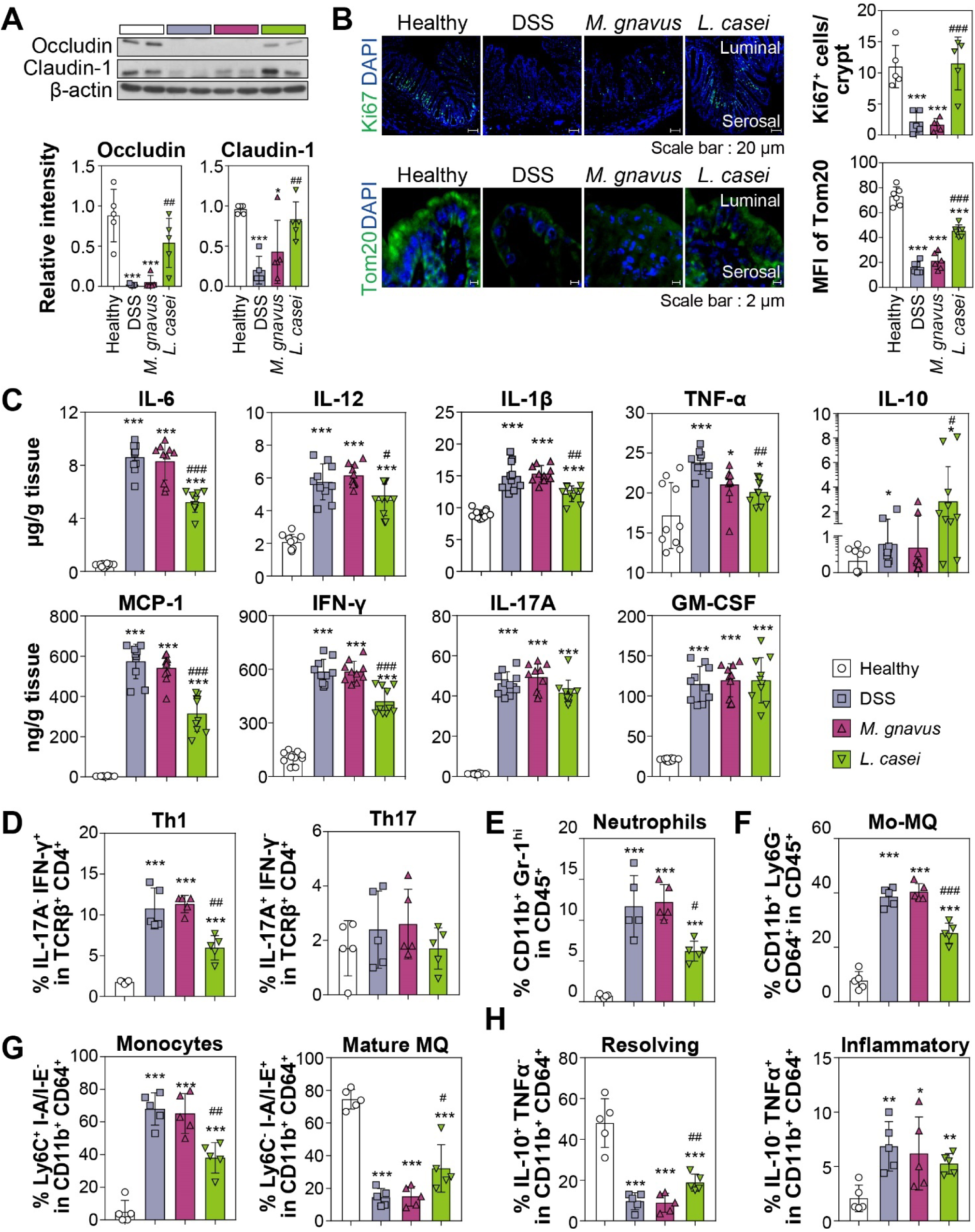
Molecular and cellular characterization of colonic responses to gut bacteria transplantation in DSS-induced colitis mice. (**A**) Western blot analysis of occludin and claudin-1 expression in distal colon tissues. Relative protein levels were quantified and presented as bar graphs (n = 4). (**B**) Immunofluorescence staining of Ki-67 (top, scale bar = 20 µm) and Tom20 (bottom, scale bar = 2 µm) in distal colon sections with quantitative analyses. (**C**) Cytokine concentrations in distal colon tissue explants measured by a cytokine bead array (n = 10). (**D** to **H**) Flow-cytometric analyses of immune cell populations in the colonic lamina propria (n = 5). Percentages of IFN-γ (Th1) and IL-17A^+^ (Th17) cells in CD45□ TCRβ□ CD4□ cells (**D**); CD11b□ Gr1^hi^ (neutrophils; **E**) and CD11b Ly6G^-^ CD64□ (monocyte-macrophages, Mo-MQ; **F**) populations in the viable CD45□ cells; Ly6C^+^ I-A/I-E^-^ (monocytes; Mo) and Ly6C^-^ I-A/I-E^+^ (mature macrophage; mature MQ) populations in CD45^+^ CD11b^+^ Ly6G^-^ CD64^+^ cells (**G**); IL-10^+^ (resolving) and TNF-α^+^ (inflammatory) macrophages in CD45^+^ CD11b^+^ Ly6G^-^ CD64^+^ cells (**H**). Data are presented as mean ± SD. Statistical significance was determined relative to the healthy or DSS control groups (\**p* < 0.05; \*\**p* < 0.01; \*\**p* < 0.001 vs. healthy control; #*p* < 0.05; ##*p* < 0.01; ^###^*p* < 0.001 vs. DSS control group).

Analysis of cytokine secretion from distal colon explants revealed that *L. casei* reduced proinflammatory cytokines, including TNF-α, IFN-γ, IL-1β, IL-6, and IL-12, while increasing the anti-inflammatory cytokine IL-10 (**Fig. 6C**). Flow cytometric analysis of the lamina propria further showed selective attenuation of IFN-γ-producing Th1 cells following *L. casei* treatment, whereas Th17 cell frequencies were unchanged (**Figs. 6D and S9A**). At the innate immune level, *L. casei* reduced the accumulation of CD11b^+^ Gr-1^hi^ neutrophils and CD11b^+^ CD64^+^ macrophages (**Figs. 6E–F, and S9B–C**). Although the proportion of TNF-α^+^macrophages was not significantly altered, *L. casei* promoted a shift from immature Ly-6C^+^ MHC-II^-^ toward more mature Ly-6C^-^ MHC-II^+^ macrophage populations, consistent with a regulated, tissue-supportive immune state (**Figs. 6G–H and S9D–E**).

To determine whether these microbial effects were dependent on inflammatory context, *M. gnavus* or *L. casei* was administered to mice in the absence of DSS. Neither bacterium altered body weight, colon length, histological features, epithelial mitochondrial mass, or cytokine profiles under steady-state conditions (**Fig. S10**), indicating that their effects become evident primarily under inflammatory stress.

Collectively, these *in vivo* findings support the epithelial-level mechanisms identified using the HoxBan system. *L. casei* consistently promotes epithelial integrity, mitochondrial competence, and balanced immune responses in both *in vitro* and *in vivo* settings, whereas *M. gnavus* induces epithelial stress programs *in vitro* but exhibits buffered effects *in vivo*, consistent with the strong homeostatic capacity of intact colonic tissues. This directional concordance underscores the utility of the HoxBan platform for mechanistic interrogation of host-microbe interactions under physiologically relevant oxic-anoxic conditions.

## Discussion

### Defining a physiologically interpretable platform for aerobic-anaerobic host-microbe interactions

Mechanistic analysis of epithelial–microbial interactions has been limited by the difficulty of sustaining oxygen-dependent epithelial cells together with strict anaerobes under defined conditions. Although the original HoxBan system enabled aerobic–anaerobic co-culture in a simple format (*29*), physicochemical parameters were not quantitatively defined, restricting interpretation of microbe-specific epithelial effects. Here, we refined the HoxBan platform to define a biologically permissive window in which epithelial integrity, metabolic competence, and oxygen architecture are preserved while allowing direct contact with anaerobic gut microbes. Incorporation of a transparent gel matrix, a mucin interface, and quantitative oxygen profiling established a reproducible vertical oxygen gradient that recapitulates the directional oxic-anoxic organization of the intestinal lumen (**Fig. 1**). Although absolute oxygen levels exceed those of the distal colon, preservation of the apical–luminal gradient provides the dominant physiological cue governing epithelial-microbial interactions (*41, 42*). Importantly, this defined operating window enables discrimination of microbe-specific epithelial programs from nonspecific cytotoxic or hypoxic stress.

Compared with Transwell-based assays or microfluidic gut-on-chip systems (*28, 32*), the optimized HoxBan platform is low-cost, scalable, and compatible with multi-omics analyses, while supporting strict anaerobes without specialized infrastructure. Although Caco-2 cells do not capture the full cellular complexity of the native colonic epithelium, they provide a stable and interpretable model for defining oxygen-interface-dependent epithelial programs (**Fig. 2**). Thus, the refined HoxBan system bridges reductionist monocultures and *in vivo* models, enabling controlled interrogation of species-dependent epithelial reprogramming under physiologically relevant oxygen partitioning.

### Divergent epithelial states induced by M. gnavus and L. casei

Application of this platform revealed strikingly divergent epithelial responses to two clinically relevant gut microbes. Integrated multi-omics analyses demonstrated that *M. gnavus* induces a coordinated epithelial stress program characterized by junctional destabilization, transcriptional reorganization, and metabolic suppression. Transcriptomic profiling revealed activation of stress-responsive signaling, including p38 MAPK signaling and chromatin-associated programs, together with repression of zinc-finger transcriptional regulators (**Fig. 3**). These changes were accompanied by reduced abundance of TJ protein abundance and suppression of mitochondrial respiration, functionally linking metabolic dysfunction to barrier destabilization.

Mitochondrial impairment is increasingly recognized as a contributor to epithelial barrier failure in UC (*43–45*). In this context, selective repression of ETCs and reduced oxygen consumption in *M. gnavus*-exposed cells provides a mechanistic link between microbial contact, epithelial energy metabolism, and TJ integrity (*46–49*) (**Fig. 2**). Consistent with its mucin-degrading capacity (*50*), *M. gnavus* upregulated pathways involved in utilization of mucin-derived substrates, suggesting reinforcement of a mucolytic metabolic state upon epithelial proximity (**Fig. S5**). Induction of pro-inflammatory mediators such as IL-6 and IL-8 further support classification of *M. gnavus* as a context-dependent pathobiont whose epithelial impact becomes pronounced when barrier resilience is compromised (**Fig. S7**), consistent with epithelial stress and innate immune activation (*51–53*). In contrast, *L. casei* elicited a restrained epithelial response characterized by preservation of junctional integrity, maintenance of mitochondrial capacity, and limited immune activation (**Figs. 2 and 3**). Co-culture enhanced ZO-1 and occludin abundance and promoted expression of mitochondrial assembly and import machinery without inducing stress-associated transcriptional programs. Proteomic signatures indicated reinforcement of mitochondrial capacity rather than acute metabolic activation (**Fig. 4**), consistent with maintenance of epithelial bioenergetic homeostasis (*52*). Cytokine outputs favored immunemodulatory and tissue-supportive profiles (**Fig. S7**), aligning with previous reports–that lactobacilli promote epithelial homeostasis through balanced metabolic support and immune restraint (*53, 54*).

Despite these opposing outcomes, both microbes induced a shared epithelial response involving downregulation of energy-intensive biosynthetic processes, as revealed by proteomic analysis (**Fig. 4**). This conserved adaptation likely reflects a general epithelial energy-conservation strategy during microbial exposure, upon which microbe-specific stress or homeostatic programs are superimposed. Differences across molecular layers further revealed temporal hierarchy: transcriptional suppression of oxidative phosphorylation preceded detectable changes in mitochondrial protein abundance following *M. gnavus* exposure, whereas *L. casei* enhanced mitochondrial assembly factors before measurable increases in respiration.

### Context-dependent in vivo relevance of epithelial programs

*In vivo* validation using a DSS-induced colitis model demonstrated that epithelial programs identified *in vitro* become physiologically relevant under inflammatory conditions. *L. casei* administration ameliorated disease severity, preserved junctional protein expression, enhanced epithelial regeneration, and increased mitochondrial mass, closely mirroring its epithelial-supportive phenotype observed in HoxBan co-cultures (**Figs. 5 and 6**). Immune profiling further clarified these effects (**Fig. 6**). While both Th1 (IFN-γ^+^) and Th17 (IL-17^+^) CD4^+^ T cells contribute to IBD pathogenesis (*55*), Immune profiling further revealed selective attenuation of Th1 responses without affecting Th17 populations (**Fig. 6**). At the innate level, excessive neutrophil infiltration is a major driver of colitis progression, whereas macrophages contribute to both inflammation and tissue repair (*56, 57*). *L. casei* reduced infiltration of CD11b^+^Gr-1^hi^ neutrophils and monocyte-macrophages populations and promoted a shift toward mature, MHC-II^+^ immunoregulatory macrophage phenotypes, indicating balanced immune modulation rather than broad immunosuppression.

Importantly, neither *L. casei* nor *M. gnavus* altered epithelial or immune parameters in healthy mice. This absence of effect likely reflects the buffering capacity of intact colonic mucosa, which masks epithelial responses that become evident only under defined oxygen-partitioned conditions or inflammatory stress. Thus, the HoxBan system and *in vivo* models provide complementary insights: the former reveals latent epithelial programs elicited by direct microbial contact, whereas the latter defines the physiological contexts in which these programs acquire functional relevance.

### Implications for microbiome-based therapeutics and functional validation

Together, these findings highlight oxygen-partitioned co-culture systems as powerful tools for uncovering microbe-specific epithelial programs relevant to intestinal disease. By enabling mechanistic distinction between stress-inducing microbes and those supporting epithelial homeostasis, the HoxBan platform provides a rational framework for therapeutic screening, biomarker discovery, and strain selection (8). Importantly, such systems offer a means for functional validation of disease-associated microbes or metabolites identified in translational studies, bridging association-based findings with mechanistic epithelial responses under defined oxygen conditions (*58*).

The modular architecture of oxygen-partitioned co-culture systems further allows stepwise incorporation of additional biological elements, enabling discrimination between direct epithelial effects and secondary influences mediated by microbial community shifts or metabolites. This flexibility also permits construction of disease-context-specific models through selective inclusion of immune components, such as Th1- or Th17-polarized T cells, facilitating interrogation of epithelial programs associated with distinct inflammatory drivers. Integration of oxygen-partitioned *in vitro* platforms with *in vivo* validation may ultimately accelerate development of microbiome-based strategies aimed at restoring epithelial–microbial homeostasis in inflammatory disorders.

## Acknowledgments

This work was supported by the Bio & Medical Technology Development Program of the National Research Foundation (NRF) of Korea grants (2021M3A9I4021431 to DWL and 2021M3A9I4023974 to KL and NJK), funded by the Ministry of Science and ICT (MSIT), Republic of Korea. This work was also partly supported by the NRF of Korea (grants RS-2025-02215096 to DWL, RS-2024-00348803 to KL, and RS-2021-NR059197 to NJK) funded by MSIT.

## Author contributions

J.E.L., T.K., S.P., J.Y.L., K.S.K., H.K., K.L., D.W.L., and N.J.K formulated the research plan, carried out experiments, analyzed and interpreted the data, and drafted the manuscript. J.E.L., T.K., S.P., K.L., D.W.L., and N.J.K participated in the design of the study and analyzed and interpreted the data. K.L., D.W.L., and N.J.K. conceived, planned, and supervised the study.

## Data and materials availability

All data supporting the findings of this study are available in the main text or the supplementary materials. Transcriptomic data have been deposited in the NCBI Sequence Read Archive (SRA) under BioProject accession number PRJNA1398586. The mass spectrometry proteomics data have been deposited to the ProteomeXchange Consortium via the PRIDE partner repository with the dataset identifier PXD072493.

