## Supplementary information for "Dual Oxygen-Partitioned Co-Culture Uncovers Microbe-Specific Epithelial Stress and Homeostatic Programs"

**The PDF file includes:**

Materials and Methods

Figures S1-S10

Table S1

Legends for Datasets 1-5

**Other Supplementary Material for this manuscript includes the following:**

Datasets 1-5

### **Supplementary Materials and Methods**

#### **Reagents and media**

Eagle's minimum essential medium (EMEM) was obtained from ATCC (Manassas, VA, USA). Fetal bovine serum (FBS), penicillin-streptomycin, trypsin-EDTA, and 0.4% trypan blue were purchased from Thermo Fisher Scientific (Waltham, MA, USA). Tryptic soy broth (TSB), de Man, Rogosa, and Sharpe (MRS) medium, and agar were obtained from BD Bioscience (Franklin Lakes, NJ, USA). Glycerol and cover glasses were purchased from Duchefa Biochemie (Haarlem, Netherlands) and Paul Marienfeld (Lauda-Königshofen, Germany), respectively. Mucin type III, Gelrite, MgCl<sub>2</sub>, Triton® X-100, and bovine serum albumin (BSA) were obtained from Sigma-Aldrich (Burlington, MA, USA), and 4% paraformaldehyde solution was purchased from Biosesang (Seongnam, South Korea). Primary antibodies against ZO-1 and occludin were obtained from Cell Signaling Technology (Danvers, MA, USA) and Invitrogen (Waltham, MA, USA), respectively. Antibodies against pan-cytokeratin (AE1/AE3+5D3, ab86734) and E-cadherin (ab40772) were purchased from Abcam (Cambridge, UK). Goat anti-rabbit IgG HRP- or fluorescein isothiocyanate (FITC)-conjugated secondary antibodies were obtained from Santa Cruz Biotechnology (Dallas, TX, USA). Unbuffered DMEM (pH 7.4), the Seahorse XFp Cell Mito Stress Test kit, XFp FluxPak, glucose, L-glutamine, and sodium pyruvate were obtained from Agilent Technologies (Santa Clara, CA, USA).

#### **Immunofluorescence microscopy**

For epithelial marker analysis, Caco-2 cells were cultured either on conventional plates or in the HoxBan system (without bacteria) for 24 h prior to fixation. Cells grown on cover glasses were fixed with 4% paraformaldehyde for 15 min, washed three times with phosphate-buffered saline (PBS), and permeabilized with 0.1% Triton X-100 for 10 min at room temperature. After washing, cells were blocked with 1% BSA in PBS containing 0.1% Tween-20 (PBS-T) for 30 min and incubated overnight at 4°C with primary antibodies against pan-cytokeratin (1:100) and E-cadherin (1:500) diluted in PBS-T. Following three PBS washes, cells were incubated with anti-rabbit-FITC-conjugated (1:500) and anti-mouse Texas Red-conjugated (1:200) secondary antibodies for 2 h at room temperature. Nuclei were counterstained with 4',6-diamidino-2-phenylindole (DAPI; 1 µg/mL). Samples were mounted and imaged using a confocal laser scanning microscope (Carl Zeiss, Germany).

To assess epithelial polarization, Caco-2 cells prepared as described above were fixed with 4% paraformaldehyde for 15 min, washed with PBS, and permeabilized with 0.1% Triton X-100 for 5 min at room temperature. After blocking with 2% BSA in PBS-T for 30 min, cells were incubated overnight at 4°C with Alexa Fluor-conjugated primary antibodies against ZO-1 (apical marker; Alexa Fluor 647, 1:400) and E-cadherin (basolateral marker; Alexa Fluor 594, 1:50). Following PBS washes, nuclei were counterstained with DAPI (1 µg/mL) for 5 min. Samples were mounted and imaged using a confocal laser scanning microscope (Carl Zeiss). Z-stack images were acquired at 0.6-µm intervals, and representative xy and xz projections were generated for analysis.

For TJ analysis, Caco-2 cells collected from the HoxBan system were fixed with 4% paraformaldehyde for 15 min and permeabilized with 0.1% Triton X-100 for 5 min. After blocking with 2% BSA for 15 min, cells were incubated overnight at 4°C with primary antibodies against ZO-1 and occludin (1:400). FITC-conjugated anti-rabbit IgG secondary antibody (1:200) was applied for 2 h at room temperature, followed by DAPI counterstaining. Samples were mounted and visualized under a fluorescence microscope (Nikon, Japan).

#### **Mitochondrial respiration analysis**

Mitochondrial respiration was assessed using the Seahorse XFp Cell Mito Stress Test (Agilent). Caco-2 cells harvested from the HoxBan system were seeded onto XFp miniplates at a density of $3 \times 10^4$ cells per

well and incubated for 24 h at 37°C in a humidified 5% CO<sub>2</sub> atmosphere. Prior to analysis, cells were washed and incubated in bicarbonate-free DMEM (pH 7.4) supplemented with 5.56 mM glucose, 2 mM L-glutamine, and 1 mM sodium pyruvate, followed by equilibration for 1 h in a non-CO<sub>2</sub> incubator. Sequential injections of oligomycin (1.5 µM), FCCP (0.5 µM), and rotenone/antimycin A (0.5 µM each) were performed using the Seahorse XFp analyzer according to the manufacturer's instructions. Oxygen consumption rate (OCR) was measured over four cycles per condition. OCR values were normalized to viable cell numbers (10<sup>4</sup> living cells) determined by trypan blue exclusion.

### **Western blotting**

After 24 h of co-culture, Caco-2 cells were washed with cold PBS and lysed in buffer containing 20 mM Tris-HCl (pH 7.5), 150 mM NaCl, 1 mM EDTA, 1 mM EGTA, 1% Triton X-100, 2.5 mM sodium pyrophosphate, 1 mM β-glycerophosphate, 1 mM Na<sub>3</sub>VO<sub>4</sub>, 1 µg/mL leupeptin, 1 mM phenylmethylsulfonyl fluoride (PMSF), and a protease inhibitor cocktail. Lysates were incubated on ice for 30 min with intermittent vortexing and centrifuged to remove insoluble debris. Protein concentrations were determined using the Bradford assay (Bio-Rad). Equal amounts of protein (20-35 µg) were separated by SDS-PAGE and transferred onto polyvinylidene fluoride (PVDF) membranes (Merck Millipore, Germany). Membranes were blocked in 5% skim milk for 2 h and incubated overnight at 4°C with primary antibodies, followed by incubation with HRP-conjugated secondary antibodies for 2 h at room temperature. Immunoreactive bands were detected using an enhanced chemiluminescence detection kit (Amersham Pharmacia Biotech). For mouse colon tissues, distal colon segments were homogenized in RIPA lysis buffer supplemented with protease and phosphatase inhibitor cocktails (GenDEPOT, TX, USA). Tissue lysates were clarified by centrifugation and processed for Western blotting as described above. Membranes were probed with primary antibodies against occludin (Abcam, UK) and claudin-1 (Santa Cruz

Biotechnology), followed by the HRP-conjugated secondary antibodies, and visualized using an enhanced chemiluminescence detection kit (Merck Millipore).

#### **Transcriptome analysis**

Total RNA was extracted from Caco-2 cells harvested after co-culture with or without gut bacteria in the HoxBan system using the RNeasy Plus Mini Kit (Qiagen) according to the manufacturer's instructions. Four independent biological replicates were prepared for each condition. RNA sequencing was performed by MacroGen Inc. (Seoul, Korea) using an Illumina sequencing platform. Sequencing libraries were generated using the TruSeq Stranded mRNA LT Sample Prep Kit (Illumina). Raw sequencing reads were evaluated for quality using FastQC (Babraham Bioinformatics) and trimmed to remove adapter sequences and low-quality bases using Trimmomatic. Processed reads were aligned to the human reference genome (GRCh38) using HISAT2.

Differential gene expression analysis was conducted using the *edgeR* package in R. Genes with zero read counts across samples were filtered out, and library size normalization was performed using the trimmed mean of M-values (TMM) method. A generalized linear model incorporating batch as a covariate was fitted, and differential expression was assessed using a likelihood ratio test. Genes exhibiting $\geq 1.5$-fold up- or downregulation with $p < 0.05$ were considered differentially expressed. Principal component analysis (PCA) was conducted on logCPM-transformed, TMM-normalized expression values after batch correction to visualize sample clustering. The first two principal components were plotted with 95% confidence ellipses. Functional enrichment analysis of differentially expressed genes (DEGs) was performed using the clusterProfiler R package with the Gene Ontology (GO) database. Immune-related transcriptional programs were further interrogated using C7 immunologic signature gene sets from the Molecular Signatures Database (MSigDB). Enrichment significance was determined using Benjamini–

Hochberg correction (adjusted $p < 0.05$).

To evaluate mitochondrial-related transcriptional changes that were not captured by enrichment-based approaches, genes associated with mitochondrial respiration, electron transport, and mitochondrial gene expression and protein handling were additionally examined using a targeted annotation overlay based on GO term pattern matching. These analyses were used for visualization and interpretation purposes and were not treated as independent enrichment tests.

#### **Proteomic analysis**

Caco-2 cells were harvested as described in the trypan blue assay. Cell pellets were resuspended in lysis buffer containing 20 mM Tris-HCl (pH 7.5), 150 mM NaCl, 1 mM Na<sub>2</sub>EDTA, 1 mM EGTA, 1% Triton X-100, 2.5 mM sodium pyrophosphate, 1 mM β-glycerophosphate, 1 mM Na<sub>3</sub>VO<sub>4</sub>, 1 μg/mL leupeptin, 1 mM PMSF, and a protease inhibitor cocktail. Lysates were incubated on ice for 30 min with intermittent vortexing and centrifuged at 10,000 × g for 10 min at 4°C. Supernatants were collected as soluble protein fractions.

Bacterial cells were recovered from the Gelrite matrix using β-agarase I (New England Biolabs). Gelrite-containing media were melted at 80°C for 30 min, cooled to 42°C, and incubated with β-agarase I for 1 h at 42°C. Bacterial pellets were collected by centrifugation at 5,000 rpm for 10 min and resuspended in 300 μL of 50 mM Tris-HCl buffer (pH 7.5) containing 150 mM NaCl. Cells were disrupted by sonification, and lysates were centrifuged at 10,000 × g for 30 min at 4°C to obtain soluble protein fractions.

For in-solution digestion, 100 μg of total protein from each sample were reduced with 5 mM tris(2-carboxyethyl)phosphine (TCEP) for 30 min at 37°C and alkylated with 6 mM iodoacetamide for 1 h in the dark. Proteins were digested overnight at 37°C with sequencing-grade trypsin at an enzyme-to-

substrate ratio of 1:25. Resulting peptides were acidified with 0.1 % formic acid (FA), desalted using C18 tips, and sequentially eluted with 40% and 60% acetonitrile (ACN) containing 0.1% FA. Eluates were dried using a centrifugal evaporator (CVE-2200, Tokyo Rikakikai Co., Ltd., Tokyo, Japan) and resuspended in 20 μL of 0.1% FA for LC-MS/MS analysis.

Peptide mixtures were analyzed on an UltiMate 3000 RSLC nano system coupled to a Q-Exactive Orbitrap HF-X mass spectrometer (Thermo Fisher Scientific), as described previously (37). Protein digests (1.5 μg) were loaded onto a trap column (75 μm × 2 cm, Acclaim PepMap 100 C18, 3 μm, 100 Å) and separated on an analytical column (75 μm × 50 cm, PepMap RSLC C18, 2 μm) at a flow rate of 0.27 μL/min. Mobile phases consisted of 0.1% FA in water (solvent A) and 0.1% (v/v) FA in ACN (solvent B). The gradient was applied as follows: 5–10% B (0–5 min), 10–35% B (5–70 min), 35–50% B (70–80 min), 50–80% B (80–85 min), hold at 80% B (85–90 min), and re-equilibration to 5% B (90–95 min). For tandem mass spectrometry, MS data were acquired in data-dependent mode with a full scan (400–2000 m/z) followed by MS/MS of the top precursor ions using normalized collision energy of 27%.

Raw MS data were processed using Proteome Discoverer (version 3.1; Thermo Fisher Scientific) and searched against proteome databases for *Homo sapiens* (UP000005640), *L. casei* ATCC 393 (NCBI Refseq assembly ID: GCF\_000829055.1), *M. gnavus* ATCC 29149 (GCF\_009831375.1), or *Mus musculus* (UniProt Proteome ID: 000000589) using the SequestHT algorithm. Precursor and fragment mass (m/z) tolerances were set to ± 10 ppm and ± 0.6 Da, respectively, allowing up to two missed cleavages. Carbamidomethylation of cysteine was set as a fixed modification, and methionine oxidation as a variable modification. Protein identifications were validated using Percolator with a false discovery rate (FDR) < 0.05.

Label-free quantification was performed based on precursor ion intensities normalized to total peptide abundance. Proteins showing ≥1.5-fold differential expression with $p < 0.05$, or uniquely detected in a

given condition, were considered differentially expressed. Functional enrichment analysis of human proteins was performed using the clusterProfiler R package with Gene Ontology (GO) terms, and C7 immunologic signature gene sets from the Molecular Signatures Database (MsigDB). Enrichment significance was determined using Benjamini–Hochberg correction (adjusted $p < 0.05$). The mass spectrometry proteomics data have been deposited to the ProteomeXchange Consortium via the PRIDE partner repository with the dataset identifier PXD072493.

### **Cytokine array assay**

Human peripheral blood mononuclear cells (hPBMCs) were obtained from ATCC and cultured at $4 \times 10^5$ cells per well in 24-well plates using phenol red-free RPMI medium supplemented with 10% FBS. After a 12 h stabilization period, hPBMCs were exposed to conditioned media collected from the apical compartment of the HoxBan co-culture system following 24 h co-culture of Caco-2 cells with *M. gnavus* or *L. casei*.

For conditioned media collection, the apical EMEM medium (1.5 mL) was replaced with phenol red-free RPMI containing 10% FBS during co-culture. Supernatants were collected after 24 h, centrifuged at $2,000 \times g$ for 10 min to remove debris, and mixed 1:1 with fresh hPBMC culture medium prior to application to hPBMCs. Control conditions included conditioned media from bacteria-free Caco-2 cultures and lipopolysaccharide (LPS, 10 $\mu\text{g/mL}$)-treated hPBMCs.

Cytokine secretion profiles were analyzed using the Proteome Profiler Human XL Cytokine Array Kit (R&D Systems, MN, USA) according to the manufacturer's instructions. Arrays were performed in duplicate. Signal intensities were quantified using HLIImage++ software (Western Vision Software). Spot intensities were normalized to the internal reference dots on each membrane, and inter-array variation was corrected based on total control signal intensity. Fold changes were calculated relative to control

conditions or expressed as normalized relative intensity values with control and LPS set to 0 and 100, respectively.

#### **DSS-induced colitis mouse model**

All animal procedures were approved by the Institutional Animal Care and Use Committee of Hallym University (approval numbers: Hallym 2021-42; Hallym 2023-62) and conducted under specific pathogen-free conditions. Male C57BL/6J mice (6 weeks old, DBL, Korea) were pretreated with 1 g/L amoxicillin (MP Biomedicals, CA, USA) in drinking water for 6 days, followed by a recovery period with normal drinking water.

Mice were orally administered $1 \times 10^9$ CFU of *M. gnavus* or *L. casei* by gavage every other day prior to and during colitis induction (**Fig. 5A**). Colitis was induced by administering 2.5% (w/v) dextran sodium sulfate (DSS; MP Biomedicals) in drinking water for 7 consecutive days. Disease activity index (DAI) was calculated as the sum of scores for body weight loss, stool consistency, and hematochezia scores, as described previously (38). Body weight loss was scored as follows: 0 (none), 1 (1-5%), 2 (5-10%), 3 (10-20%), and 4 (>20%). Stool consistency was scored as 0 (firm), 1 (loose), and 4 (diarrhea). Bleeding was scored as 0 (none), 1 (hemocult-positive), 2 (visible blood), and 4 (gross bleeding).

#### **Histopathology and Immunofluorescence**

Distal colon tissues were fixed in 4% paraformaldehyde, dehydrated, embedded in paraffin, and sectioned at 5 μm thickness. Sections were stained with hematoxylin and eosin (H&E) for histological evaluation or with periodic acid-Schiff (PAS)–Alcian blue for visualization of goblet cells. H&E-stained sections were scored in a blinded manner for epithelial damage (0, normal; 1, hyperproliferation; 2, <50% crypt loss; 3, >50% crypt loss; 4, complete crypt loss; 5, ulceration) and inflammatory cells infiltration (0, none; 1, mild;

2, moderate; 3, severe), with a maximum combined score of 11.

For immunofluorescence analysis, tissue sections were permeabilized in PBS containing 0.2% Triton X-100 and incubated with primary antibodies against Ki-67 (Thermo Fisher Scientific) or Tom20 (Cell Signaling Technology), followed by Alexa Fluor 488–conjugated secondary antibodies. Nuclei were counterstained with DAPI using ProLong™ Diamond Antifade Mountant (Thermo Fisher Scientific). Images were acquired using an Axio Scope 5 fluorescence microscope (Carl Zeiss). Ki-67+ cells per crypt and mean fluorescence intensity (MFI) of Tom20 within defined regions of interest in distal colon tissues were quantified using ZEN microscopy software (Carl Zeiss) according to the manufacturer’s instructions.

##### **Enzyme-linked immunosorbent assay (ELISA)**

Mouse fecal samples were collected five days after DSS administration. Individual fecal pellets were homogenized in 200 µL PBS and centrifuged at 14,000 rpm for 10 min to remove debris. Lipocalin-2 and calprotectin levels were quantified using Mouse Lipocalin2/NGAL and S100A8/S100A9 (calprotectin) ELISA Kits (R&D Systems) according to the manufacturer’s instructions. Cytokine concentrations were normalized to fecal pellet weight.

##### **Fecal DNA extraction and 16S rRNA amplicon sequencing**

Fecal DNA was extracted using the QIAamp PowerFecal Pro DNA Kit (QIAGEN). Amplicon libraries were prepared using 12.5 ng DNA per sample following Illumina’s 16S metagenomic library preparation protocol. The V1-V3 hypervariable regions were amplified using primers 27F (5’-AGAGTTTGATCCTGGCTCAG-3’) and 534R (5’-ATTACCGCGGCTGCTGG-3’) and sequenced on an Illumina MiSeq platform.

Sequencing reads were trimmed using fastp (v0.23.276), merged as described previously (39), and

processed with DADA2 to generate amplicon sequence variants (ASVs). Taxonomic assignment was performed using the SILVA database (v138.1) within QIIME 2 (v2022.2). Alpha- and beta-diversity analyses were conducted using phyloseq (40), vegan (41), and ggplot2 R packages. ASVs with relative abundance below 0.25% across all samples were excluded prior to normalization. Differentially abundant taxa were identified using linear discriminant analysis effect size (LEfSe).

#### **Colon tissue cytokine measurement**

Distal colon segments (~10 mm) were excised, rinsed with PBS, and incubated in 500 μL Iscove's modified Dulbecco's medium (IMDM) supplemented with 10% FBS at 37°C under 5% CO<sub>2</sub>. After 24 h, supernatants were centrifuged at 14,000 rpm for 10 min to remove debris and analyzed using the LEGENDplex Mouse Inflammation Panel (13-plex; BioLegend) according to the manufacturer's protocol. Cytokine concentrations were normalized to tissue weight and quantified using a FACSCalibur flow cytometer (BD Biosciences).

#### **Flow Cytometry**

Single-cell suspensions from colonic lamina propria were prepared using the Mouse Lamina Propria Dissociation Kit (Miltenyi Biotech) and a gentleMACS Dissociator (Miltenyi Biotech). Cells were blocked with Mouse BD Fc Block (BD Biosciences) and stained with fluorophore-conjugated antibodies against CD11b (M1/70), Ly-6G (1A8), CD64 (X54-5/7.1), CD4 (RM4-5), and I-A/I-E (M5114.15.2) (BD Biosciences), or TCR-β (H57-597) and Ly-6C (HK1.4) (eBioscience).

For intracellular cytokine staining, cells were stimulated with 50 ng/mL phorbol 12-myristate 13-acetate (PMA) and 1 μL/mL ionomycin (Sigma) in the presence of Golgi-Stop and Golgi-Plug reagents (BD Biosciences) for 6 h at 37°C. Cells were then fixed, permeabilized, and stained for IFN-γ (XMM1.2),

TNF-α (MP6-XT22) (BD Biosciences), IL-17A (eBio17B7), and IL-10 (JES5-16E3) (eBiosciences). Data were acquired using a FACSCanto II cytometer (BD Biosciences) and analyzed with FlowJo V10 software (BD Biosciences).

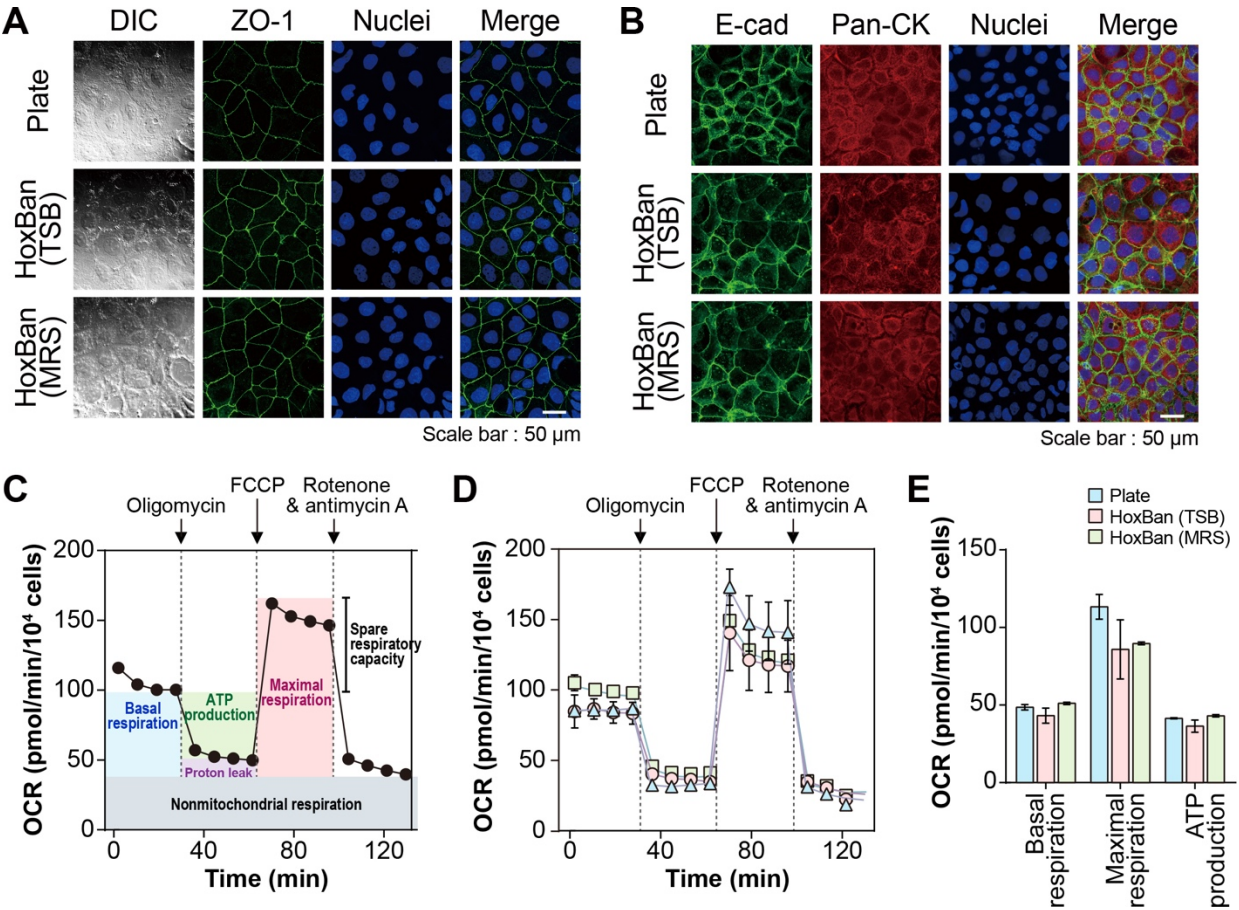

**Figure S1. Structural and functional validation of epithelial cells cultured in the HoxBan system in the absence of bacteria.** (A) Representative differential interference contrast (DIC) and confocal immunofluorescence images of Caco-2 cells cultured under conventional plate conditions or in the HoxBan system using TSB or MRS medium in the absence of bacteria. Cells were stained for the TJ protein ZO-1 (green), and nuclei were counterstained with DAPI (blue). (B) Immunofluorescence staining of the adherens-junction marker E-cadherin (E-cad, green) and epithelial marker pan-cytokeratin (Pan-CK, red) in Caco-2 cells cultured under Plate and HoxBan conditions. Nuclei were counterstained with DAPI (blue). (C) Schematic overview of the Seahorse XFp mitochondrial stress-test protocol. Sequential injections of oligomycin (1.5 μM), FCCP (0.5 μM), and rotenone/antimycin A (0.5 μM) were used to derive mitochondrial respiration parameters. (D) Representative oxygen-consumption-rate (OCR) profiles of Caco-2 cells cultured under Plate, HoxBan (TSB), and HoxBan (MRS) conditions in the absence of bacteria. (E) Quantification of basal respiration, maximal respiration, and ATP-linked respiration derived from OCR measurements is shown in (D). All experiments were performed after 24 h of culture. OCR values were normalized to cell number and are presented as mean ± SD from three independent experiments. Scale bars = 50 μm.

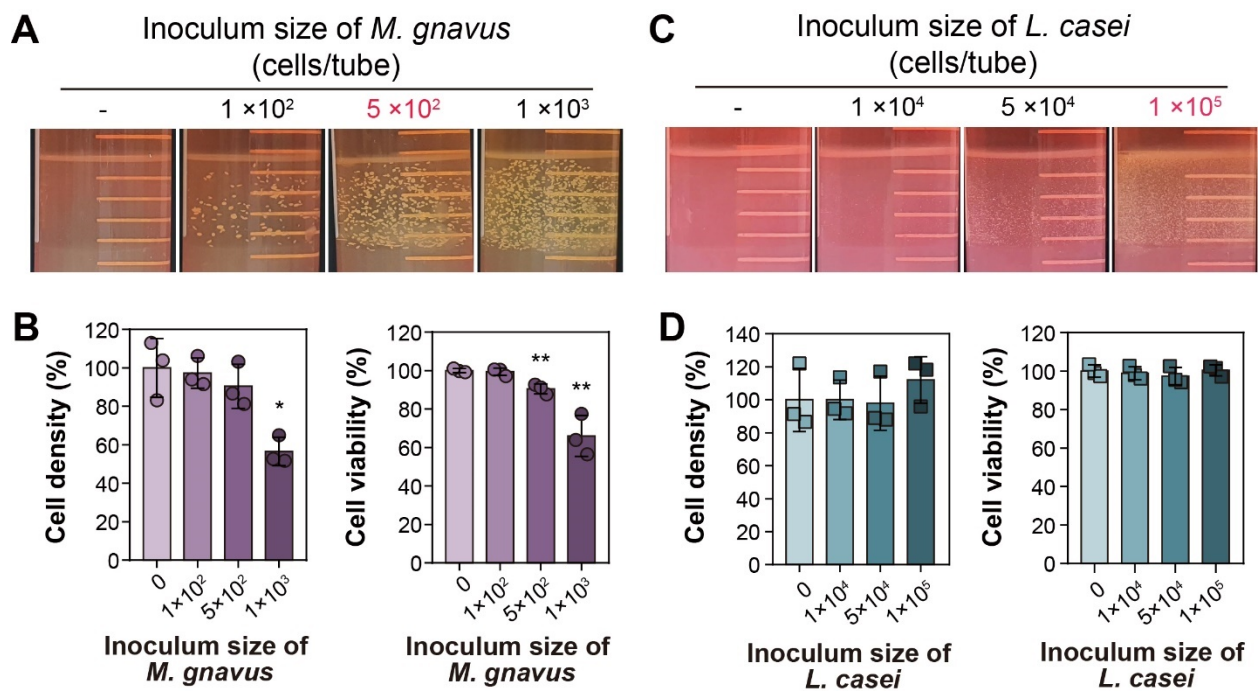

**Figure S2. Microbial growth and epithelial response in the modified HoxBan co-culture system.** (A) Inoculum-dependent growth patterns of *M. gnavus* on the anaerobic layer of the HoxBan system. (B) Density and viability of Caco-2 cells after 24 h of co-culture with *M. gnavus*. (C) Inoculum-dependent growth patterns of *L. casei* on the anaerobic layer of the HoxBan system. (D) Density and viability of Caco-2 cells after 24 h of co-culture with *L. casei*. Gut bacteria were inoculated at the indicated cell densities per tube and co-cultured with Caco-2 cells for 24 h. Bacterial growth was evaluated visually, and epithelial cell density and viability were assessed by Trypan Blue staining. Quantitative data are presented as mean ± SD (n = 3) relative to the group. Statistical significance was determined using a *t*-test versus the without-bacteria group (\**p* < 0.05; \*\**p* < 0.01).

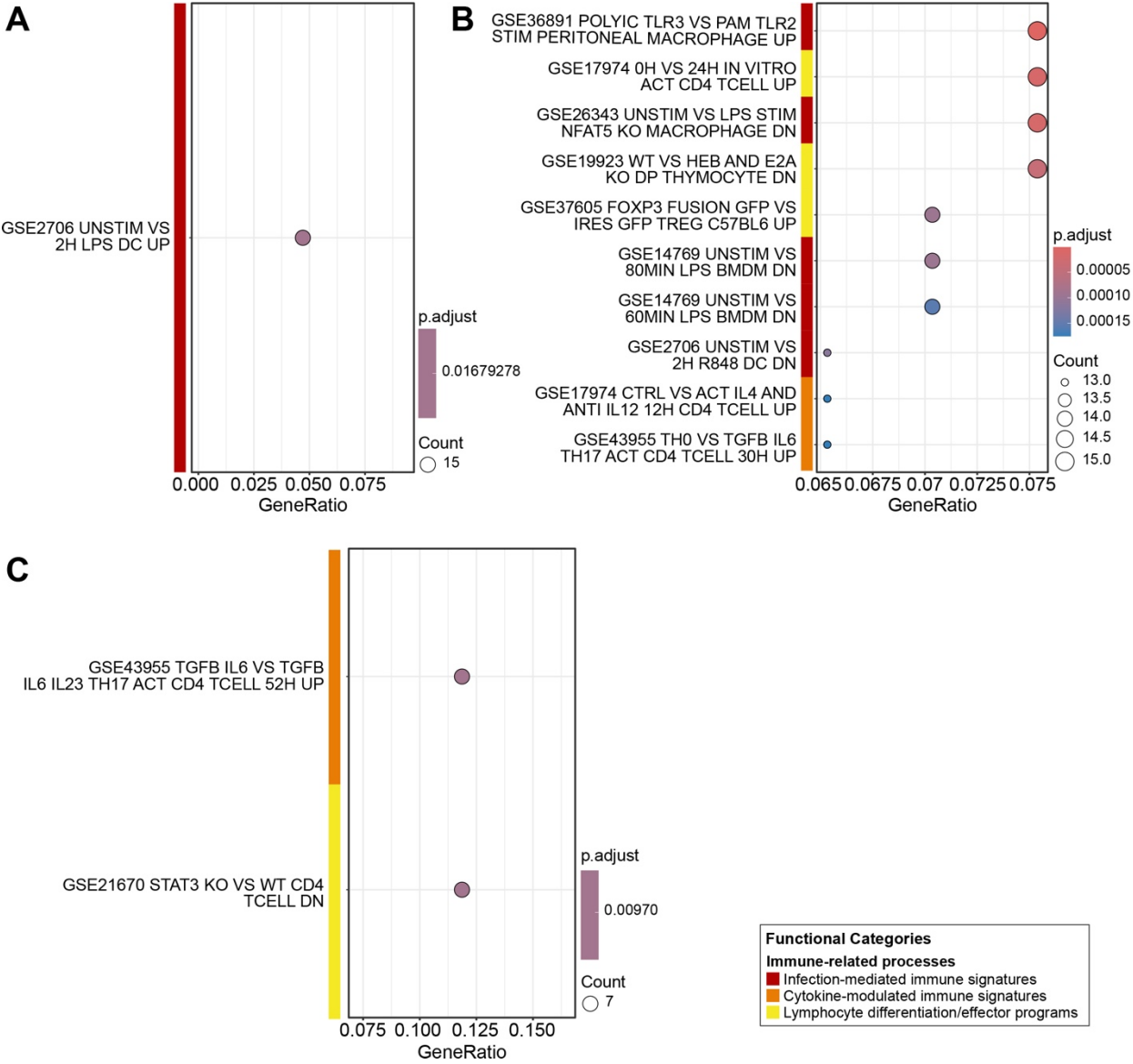

**Figure S3. Over-represented immune-related gene signatures in Caco-2 cells after co-culture with** **gut bacteria. (A and B)** Dot plots of enriched immune-related signature terms derived from downregulated (A) and upregulated (B) genes in Caco-2 cells after 24 h co-culture with *M. gnavus*. (C) Dot plots of enriched immune-related gene signatures terms derived from upregulated genes in Caco-2 cells after co-culture with *L. casei*. Enrichment analysis was performed against curated immune-related gene signature collections from MSigDB(C7). Dot plots show significantly enriched (adjusted $p < 0.05$), ranked by gene ratio. For each node, the dot size represents the number of overlapping genes, and the color indicates the adjusted $p$-value. Enriched signatures were grouped into three functional categories: infection-mediated immune signatures (red), cytokine-modulated immune signatures (orange), and lymphocyte differentiation or effector programs (yellow).

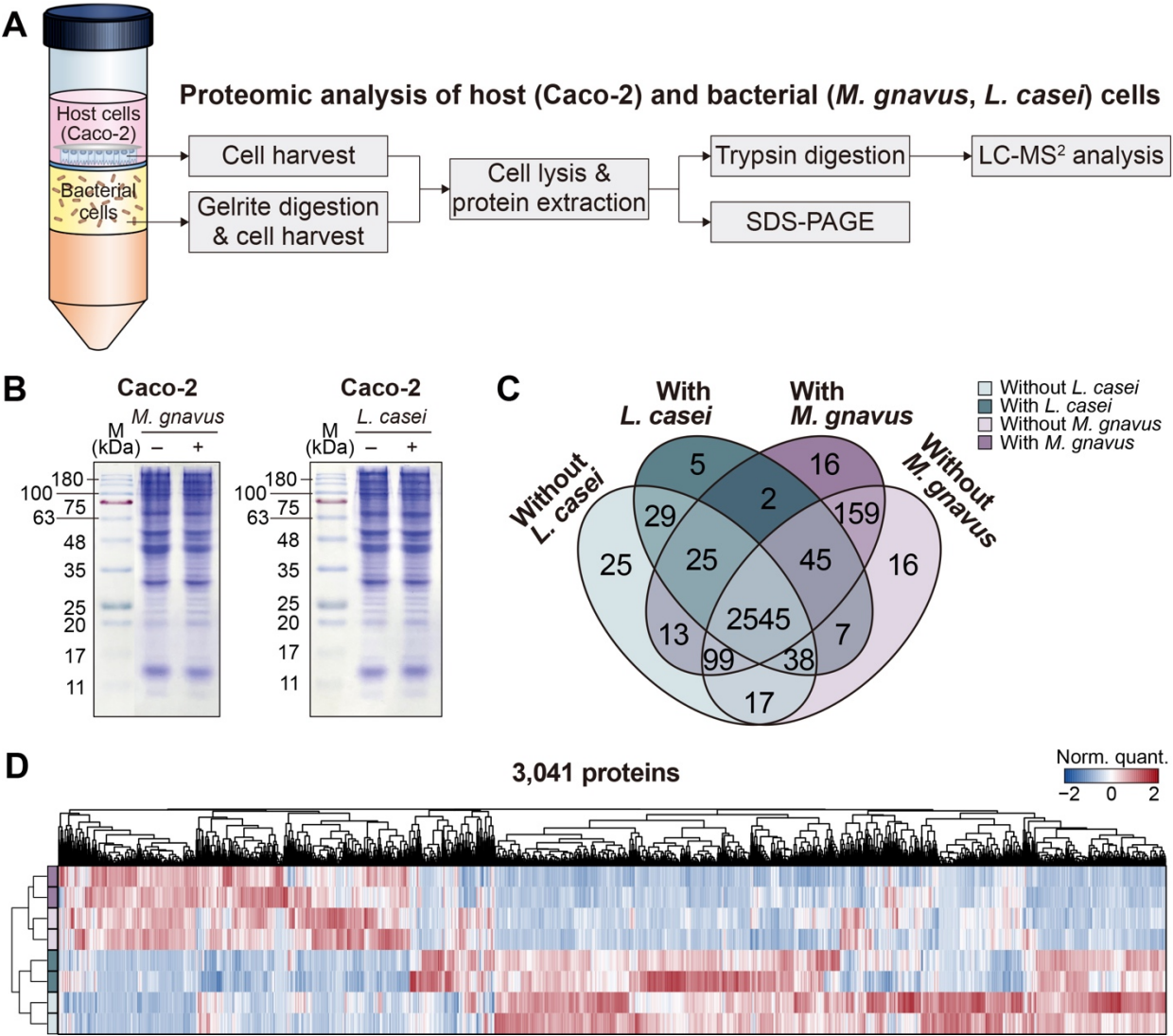

**Figure S4. Proteomic profiling of epithelial cells after co-culture in the HoxBan system.** (A) Schematic overview of the workflow used for proteomic analysis of host (Caco-2) and bacterial (*M. gnnavus* and *L. casei*) cells following co-culture in the HoxBan system. Host and bacterial compartments were harvested separately, followed by protein extraction, enzymatic digestion, and LC–MS/MS analysis. (B) SDS-PAGE analysis of soluble protein extracts from Caco-2 cells cultured in the absence or presence of *M. gnnavus* and *L. casei*, illustrating overall proteome complexity and comparable protein loading across conditions. (C) Venn diagram showing the overlap of the human proteins identified in Caco-2 cells cultured without bacteria or co-cultured with *M. gnnavus* or *L. casei*. A total of 3,041 proteins were detected across all conditions. (D) Unsupervised hierarchical clustering of normalized protein abundances in Caco-2 cells across the indicated conditions. Rows represent proteins and columns represent individual samples. Protein abundance values were Z-score-normalized, with red and blue indicating relatively higher and lower expression levels, respectively.

325

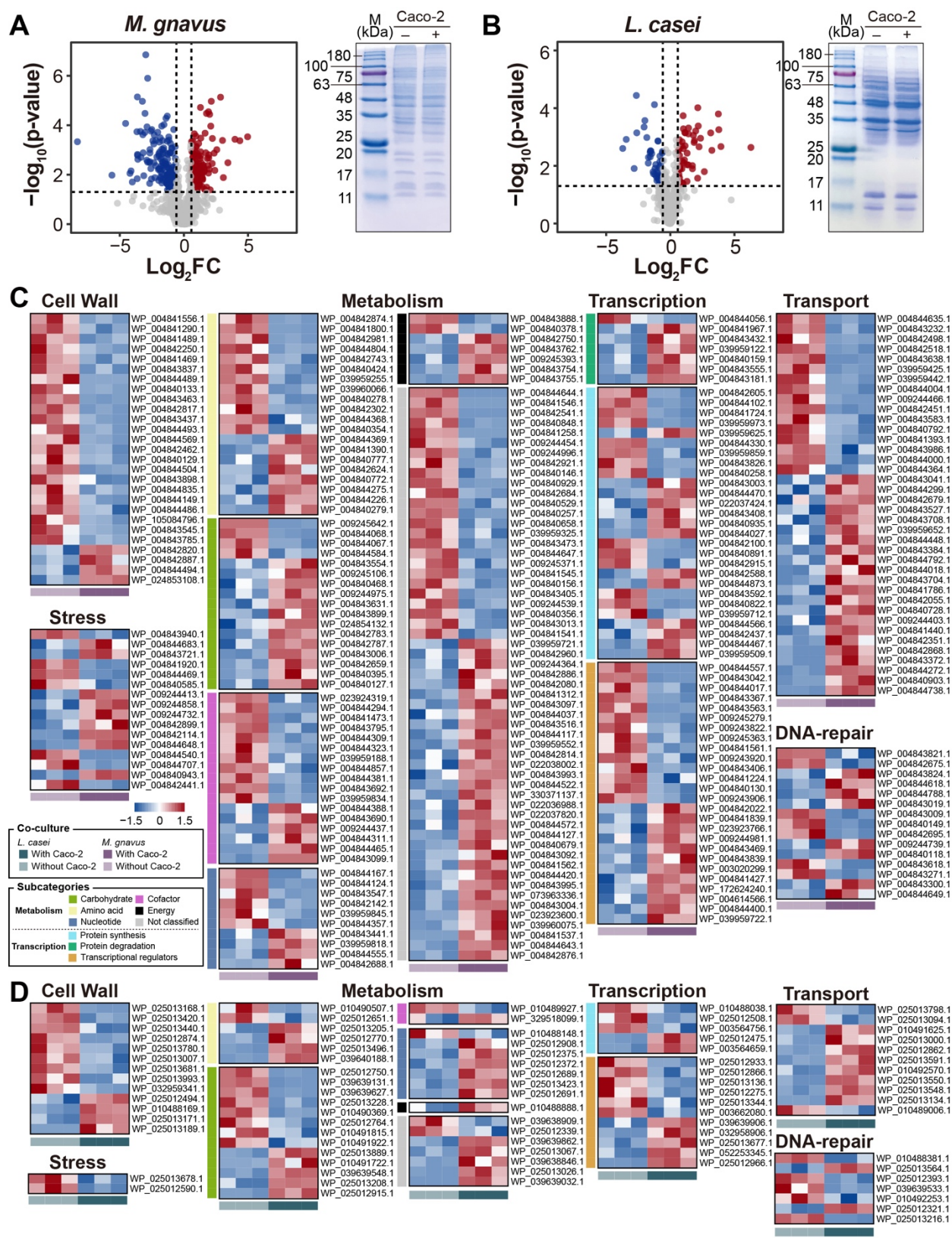

**Figure S5. Proteomic remodeling of gut bacteria co-culture with Caco-2.** (A and B) Volcano plots showing differentially expressed proteins in *M. gnavus* (A) and *L. casei* (B) following co-culture with

Caco-2 cells in the HoxBan system. Proteins with significant upregulation ($\log_2(\text{FC}) \geq \log_2(1.5)$, $p < 0.05$) and downregulation ($\log_2(\text{FC}) \leq -\log_2(1.5)$, $p < 0.05$) are shown in red and blue, respectively; non-significant proteins are shown in grey. Representative SDS–PAGE profiles of bacterial soluble proteomes recovered with (+) or without (–) Caco-2 co-culture are shown to the right. **(C and D)** Heatmaps showing differentially expressed bacterial proteins in *M. gnavus* **(C)** and *L. casei* **(D)** following co-culture with Caco-2 cells, relative to bacteria-only controls. Proteins are grouped according to major functional categories, including cell wall, transport, metabolism, transcription, stress, DNA repair, and cell division. Each value represents normalized protein abundances, in which red and blue represent high and low protein expressions, respectively.

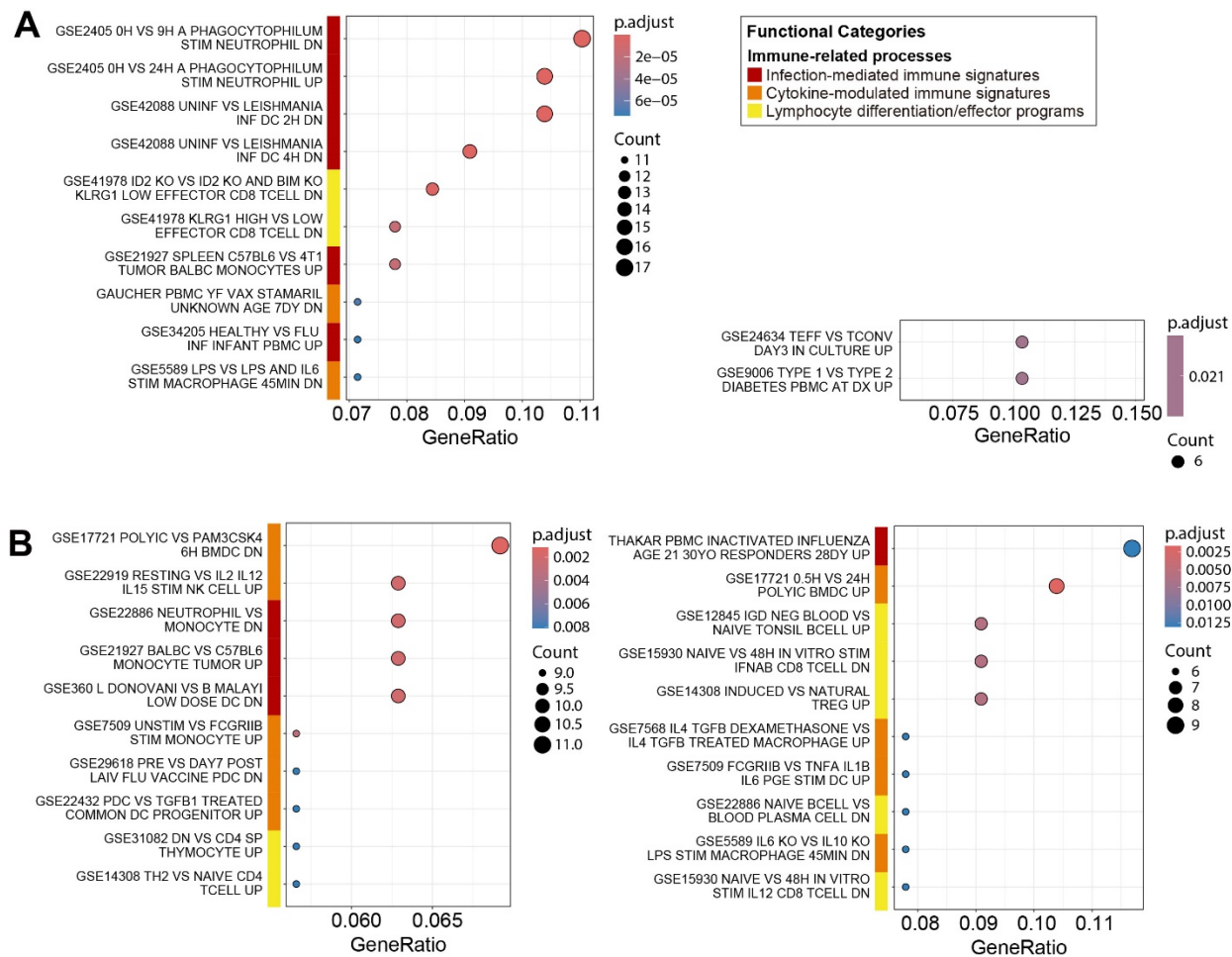

**Figure S6. Over-represented immune-related protein signatures in Caco-2 after co-culture with gut** **bacteria. (A and B) Dot plots of enriched immune-related signature terms derived from downregulated** **(left) and upregulated (right) proteins after 24 h co-culture with *M. gnavus* (A) and *L. casei* (B).** **Enrichment analysis was performed against curated immune-related gene signature collections from** **MSigDB(C7). Dot plots show significantly enriched top 10 GO terms (adjusted $p < 0.05$), ranked by gene** **ratio. For each node, size and color represent the number of genes and adjusted $p$-values, respectively.** **Individual terms are categorized into three categories: infection-mediated immune signatures (red),** **cytokine-modulated immune signatures (orange), and lymphocyte differentiation/effector programs** **(yellow).**

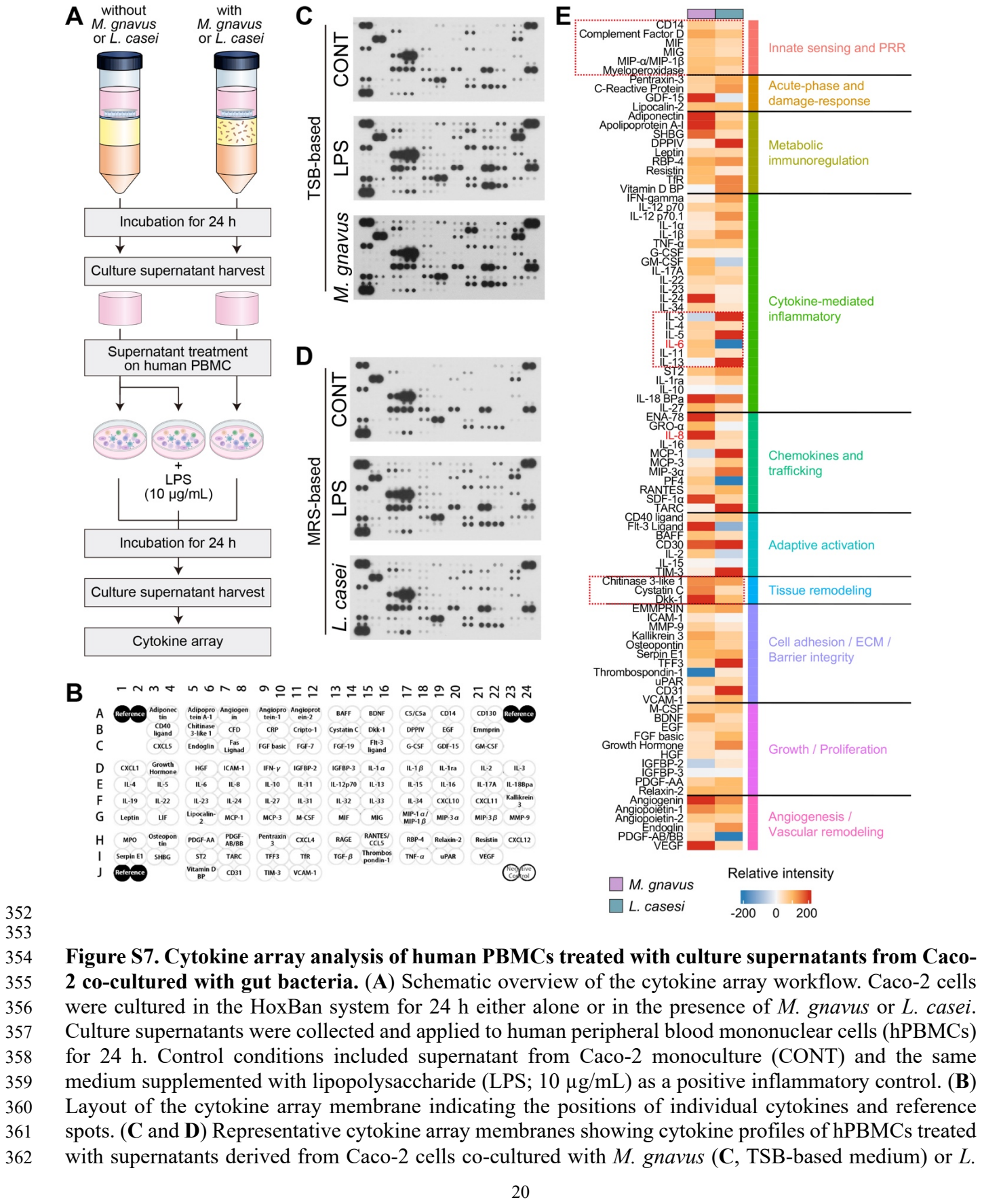

Figure S7. Cytokine array analysis of human PBMCs treated with culture supernatants from Caco-2 co-cultured with gut bacteria. (A) Schematic overview of the cytokine array workflow. Caco-2 cells were cultured in the HoxBan system for 24 h either alone or in the presence of M. gnavus or L. casei. Culture supernatants were collected and applied to human peripheral blood mononuclear cells (hPBMCs) for 24 h. Control conditions included supernatant from Caco-2 monoculture (CONT) and the same medium supplemented with lipopolysaccharide (LPS; 10 µg/mL) as a positive inflammatory control. (B) Layout of the cytokine array membrane indicating the positions of individual cytokines and reference spots. (C and D) Representative cytokine array membranes showing cytokine profiles of hPBMCs treated with supernatants derived from Caco-2 cells co-cultured with M. gnavus (C, TSB-based medium) or L.

363 *casei* (**D**, MRS-based medium), alongside corresponding control and LPS-treated conditions. (**E**) Heatmap
364 summarizing relative cytokine intensities across conditions. Signal intensities were normalized to the
365 Caco-2 monoculture control (set to 0) and expressed relative to the LPS (10 µg/mL)-treated condition (set
366 to 100). Cytokines are grouped by functional categories. Dashed boxes highlight cytokines showing
367 prominent differential induction between bacterial conditions.

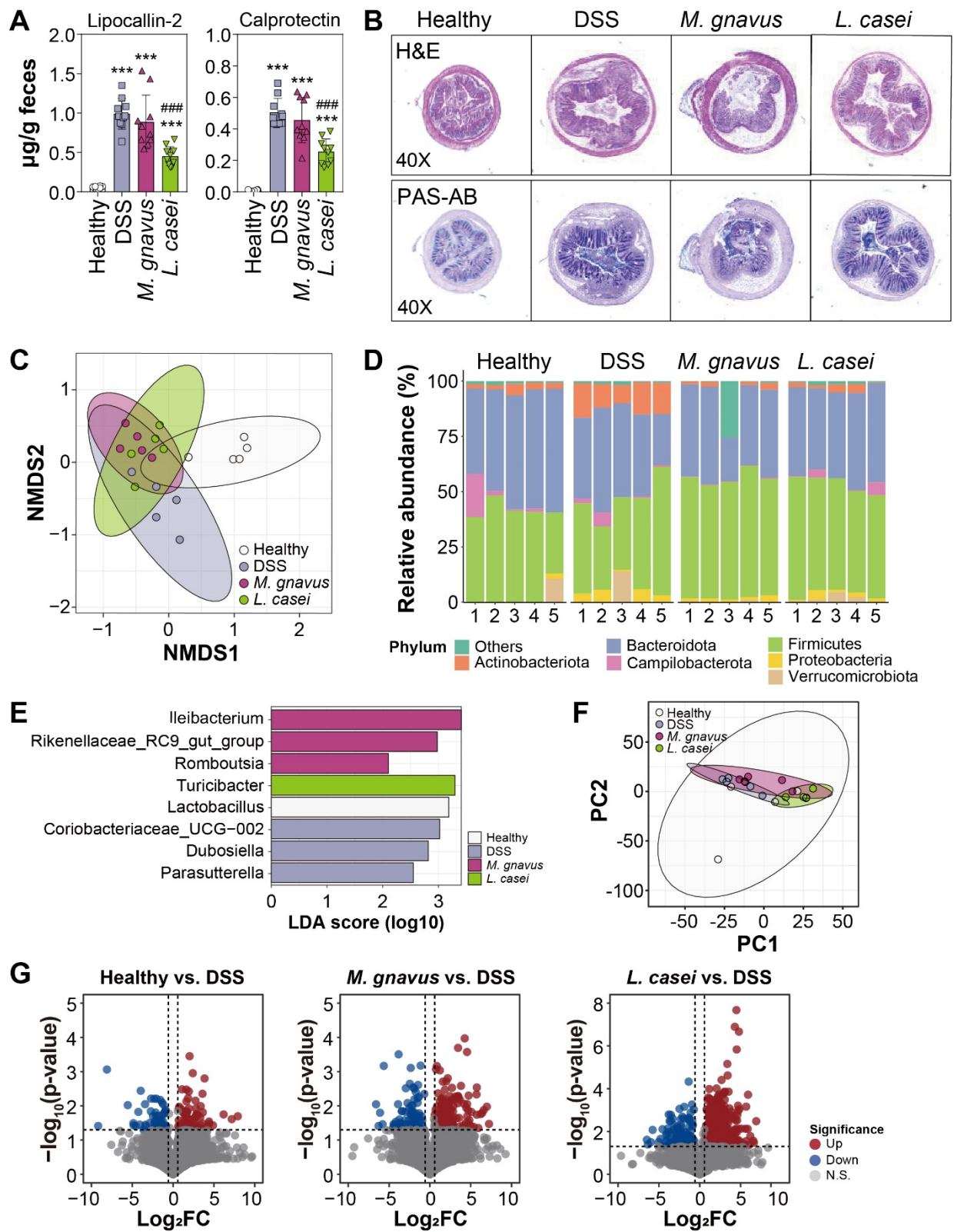

**Figure S8. Fecal inflammatory markers, gut microbiota composition, and colonic proteomic remodeling in DSS-induced colitis. (A) Fecal levels of lipocalin-2 and calprotectin, measured by ELISA**

as indicators of intestinal inflammation (n = 10 per group). **(B)** Representative distal colon sections stained with hematoxylin and eosin (H&E) and PAS-Alcian blue (PAS-AB) to evaluate mucosal architecture and goblet-cell depletion. Images are shown at $\times 40$ magnification. **(C)** Non-metric multidimensional scaling (NMDS) analysis based on Bray–Curtis distances, showing β-diversity of gut microbiota across Healthy, DSS, *M. gnavus*-, and *L. casei*-treated groups. Shaded ellipses indicate 95% confidence intervals. **(D)** Phylum-level relative abundance of gut microbiota. “Others” represents taxa comprising $< 1\%$ of total sequences. **(E)** Linear discriminant analysis (LDA) effect size (LEfSe) identifying taxa differentially enriched among experimental groups (LDA score $> 2$). **(F)** Principal component analysis (PCA) of proteomic profiles from mouse colon tissues in Healthy, DSS-induced, *M. gnavus*, and *L. casei* groups. **(G)** Volcano plots showing differentially expressed proteins in colon tissues from each group compared with DSS controls. Significantly upregulated [$\log_2(\text{FC}) > \log_2(1.5)$ in normalized protein abundance, $p <$ $0.05$] and downregulated proteins [$\log_2(\text{FC}) < -\log_2(1.5)$, $p < 0.05$] are shown in red and blue, respectively; non-significant (N.S.) proteins are shown in grey.

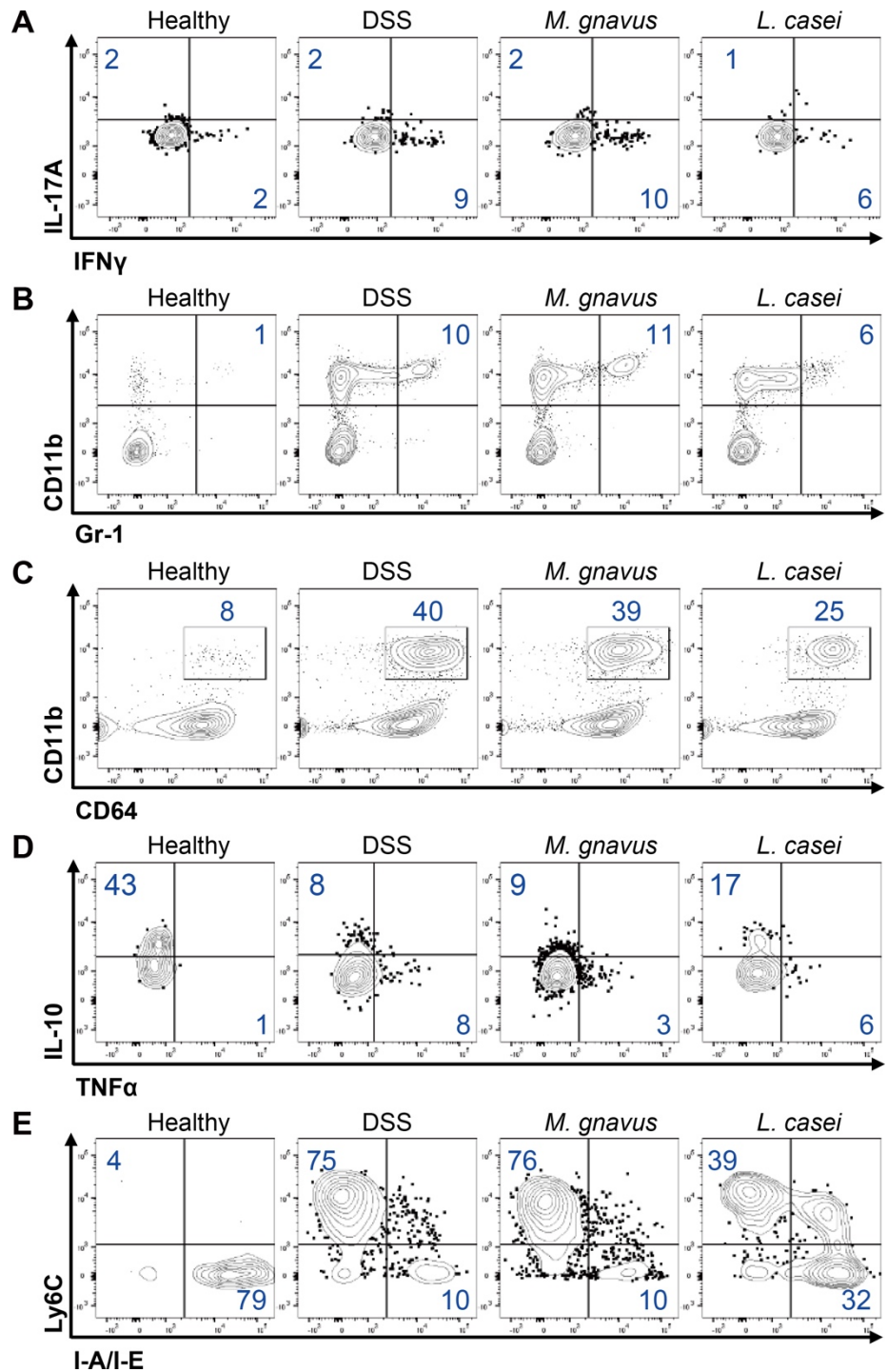

**Figure S9. Flow cytometric profiling of immune cell populations in the colonic lamina propria during DSS-induced colitis.** (A) Representative flow cytometry profiles of CD4 T cells producing IFN-γ and IL-17A are shown in the viable CD45<sup>+</sup> TCR-β<sup>+</sup> CD4<sup>+</sup> gate. (B and C) Prevalence of neutrophils (CD11b<sup>+</sup> Gr1<sup>hi</sup>) and macrophages (CD11b<sup>+</sup> CD64<sup>+</sup>) is shown in the viable CD45<sup>+</sup> gate. (D and E) Representative FACS profiles of macrophages expressing the indicated surface markers (D) or cytokines (E) are shown in the viable CD45<sup>+</sup> CD11b<sup>+</sup> CD64<sup>+</sup> gate (n = 5).

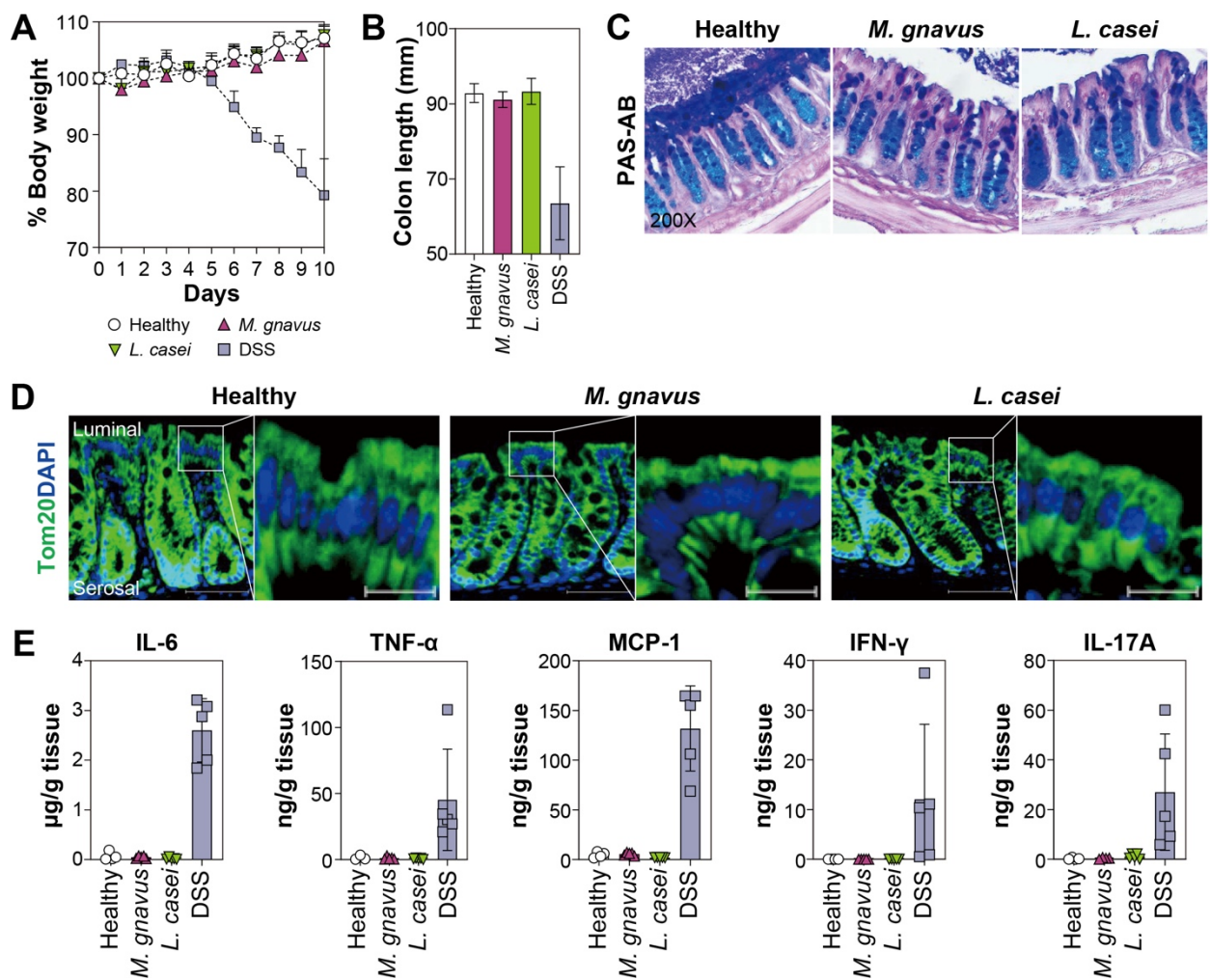

**Figure S10. *In vivo* effects of *M. gnnavus* and *L. casei* administration under steady-state (non-inflamed) conditions.** (A) Body weight changes are monitored daily following oral administration of *M. gnavus* or *L. casei* in the absence of DSS treatment. (B) Colon length was measured 10 days after bacterial administration. (C) Representative histological images of distal colon sections stained with PAS–Alcian blue, showing preserved mucosal architecture and goblet cell distribution (200×). (D) Representative immunofluorescence images of distal colon sections stained for the mitochondrial marker Tom20. (E) Cytokine concentrations in distal colon tissue explants were measured by cytokine bead array. No significant differences were observed among Healthy, *M. gnnavus*–treated, and *L. casei*–treated groups across all measured parameters.

412 **Table S1. Summary of RNA-sequencing raw data and mapping statistics.** The table summarizes
413 sequencing quality and mapping metrics for each RNA-seq library. Raw data statistics include total read
414 length (bp), total reads, GC content (%), and sequencing quality scores (Q20, Q30). Mapping statistics
415 indicate the number and proportion of mapped and unmapped reads for each RNA-sequencing library.
416 These metrics collectively ensure data quality for downstream analysis.
417

| Samples | Raw data statistics |  |  |  |  | Mapped data statistics |  |  |  |
| --- | --- | --- | --- | --- | --- | --- | --- | --- | --- |
|  | Total read bases | Total reads | GC (%) | Q20 (%) | Q30 (%) | No. of mapped reads | (%) | No. of unmapped reads | (%) |
| w/o. *M. gnavus* 1 | 7,267,108,570 | 71,951,570 | 50.29 | 98.24 | 94.95 | 70,284,498 | 99.16 | 593,124 | 0.84 |
| w/o. *M. gnavus* 2 | 7,918,570,084 | 78,401,684 | 50.33 | 98.17 | 94.87 | 76,403,722 | 99.12 | 680,630 | 0.88 |
| w/o. *M. gnavus* 3 | 8,410,991,544 | 83,277,144 | 50.10 | 98.58 | 95.94 | 81,306,427 | 99.04 | 790,161 | 0.96 |
| w/o. *M. gnavus* 4 | 8,750,506,276 | 86,638,676 | 49.99 | 98.53 | 95.87 | 84,466,017 | 99.03 | 823,137 | 0.97 |
| w/. *M. gnavus* 1 | 6,922,773,310 | 68,542,310 | 50.41 | 98.14 | 94.83 | 66,725,065 | 99.09 | 616,089 | 0.91 |
| w/. *M. gnavus* 2 | 7,436,367,400 | 73,627,400 | 51.46 | 98.27 | 95.06 | 71,802,340 | 99.09 | 661,076 | 0.91 |
| w/. *M. gnavus* 3 | 8,436,971,370 | 83,534,370 | 49.84 | 98.48 | 95.73 | 81,280,935 | 98.88 | 922,727 | 1.12 |
| w/. *M. gnavus* 4 | 8,535,420,110 | 84,509,110 | 50.50 | 98.55 | 95.91 | 82,480,544 | 99.06 | 785,832 | 0.94 |
| w/o. *L. casei* 1 | 8,493,405,524 | 84,093,124 | 49.14 | 98.53 | 95.37 | 82,504,783 | 99.21 | 659,083 | 0.79 |
| w/o. *L. casei* 2 | 7,182,007,788 | 71,108,988 | 50.46 | 98.06 | 94.63 | 69,229,510 | 99.12 | 613,340 | 0.88 |
| w/o. *L. casei* 3 | 9,013,546,232 | 89,243,032 | 50.93 | 98.50 | 95.73 | 87,165,167 | 99.16 | 740,083 | 0.84 |
| w/o. *L. casei* 4 | 7,190,733,178 | 71,195,378 | 49.13 | 99.01 | 96.89 | 69,865,453 | 98.79 | 852,719 | 1.21 |
| w/. *L. casei* 1 | 7,223,893,902 | 71,523,702 | 49.11 | 98.40 | 95.19 | 69,968,304 | 99.05 | 674,600 | 0.95 |
| w/. *L. casei* 2 | 6,891,851,150 | 68,236,150 | 50.49 | 98.05 | 94.55 | 66,423,368 | 99.14 | 578,856 | 0.86 |
| w/. *L. casei* 3 | 7,855,195,816 | 77,774,216 | 50.33 | 98.51 | 95.78 | 75,908,760 | 99.10 | 687,430 | 0.90 |
| w/. *L. casei* 4 | 7,053,647,898 | 69,838,098 | 49.56 | 99.09 | 97.11 | 68,602,676 | 98.81 | 827,536 | 1.19 |

418
419

**DESCRIPTION OF THE SUPPLEMENTAL TABLES (provided as a separate file)**

**Dataset 1. Transcriptomic profiles of Caco-2 cells co-cultured with *M. gnavus* or *L. casei*.** Fold changes and statistics for the transcriptome in Caco-2 cells under each co-culture condition.

**Dataset 2. Over-representation analysis of differentially expressed genes in Caco-2 cells during co-culture.** GO terms or C7 gene sets (immunologic signature) from the molecular signatures database (MSigDB) were significantly enriched among up- or downregulated proteins in Caco-2 cells co-cultured with *M. gnavus* or *L. casei*.

**Dataset 3. Proteomic profiles of Caco-2 cells, bacteria, and colon tissues.** Differentially expressed proteins were identified in Caco-2 cells and bacterial partners during co-culture, and in colon tissues from healthy, DSS, and bacteria-treated mice.

**Dataset 4. Over-representation analysis of differentially expressed proteins in Caco-2 cells during co-culture.** GO Biological Process terms or C7 gene sets (immunologic signature) from the molecular signatures database (MSigDB) were significantly enriched among up- or downregulated proteins in Caco-2 cells co-cultured with *M. gnavus* or *L. casei*.

**Dataset 5. Relative intensity of cytokine responses in hPBMC treated with culture supernatant from Caco-2 co-cultured with gut bacteria.** Fold-changes (FC) and *p*-value relative to bacteria-free controls (CONT), LPS reference comparisons, and normalized cytokine intensities (%; control = 0 and LPS = 100).
